# Triangulating evidence to detect signatures of stabilizing selection acting on protein levels in humans

**DOI:** 10.64898/2026.10.16.759432

**Authors:** Mihaela-Diana Zanoaga, Théo Cavinato, Thibault Latrille, Orsolya Liska, Tabea Schoeler, Balázs Papp, Zoltán Kutalik

**Affiliations:** Department of Computational Biology, University of Lausanne, Lausanne, Switzerland; SIB Swiss Institute of Bioinformatics, Lausanne, Switzerland; Unisanté, Lausanne, Switzerland; Biological Research Centre & HCEMM, Szeged, Hungary

## Abstract

Stabilizing selection is thought to maintain quantitative traits near evolutionary optima, yet the extent to which it constrains human molecular phenotypes remains largely unknown. Here, we investigate stabilizing selection on 2’923 plasma proteins by triangulating complementary signatures operating across evolutionary timescales in up to 54’000 UK Biobank participants. We contrasted genetic signatures reflecting selective constraint accumulated over generations with phenotypic evidence that deviations in protein abundance are associated with reduced contemporary fitness-related outcomes. Historical selection was assessed from the frequency-dependent architecture of protein levels using genome-wide common variants with LDpred2 and a newly developed rare loss-of-function burden framework. Contemporary evidence was obtained from quadratic associations between protein abundance and three fitness proxies: reproductive success, somatic health, and parental longevity. Among significant estimates, 100% and 92.4% supported stabilizing selection in the common- and rare-variant-based analysis, respectively, compared to 99.8% for somatic health, 97.7% for parental longevity, and 89.3% for reproductive success. Despite capturing distinct timescales and aspects of selection, the approaches showed convergence. Rare-variant-based estimates showed concordance with health-score-derived coefficients (*ρ* = 0.36, *p* = 5.2 *×* 10*^−^*^6^) and with independent measures of evolutionary constraint, including protein heritability (*ρ* = 0.20, *p* = 5.2 *×* 10*^−^*^5^), loss-of-function intolerance (*ρ* = *−*0.19, *p* = 6.1 *×* 10*^−^*^5^), and cross-species transcript evolutionary rate (*ρ* = 0.15, *p* = 2.4 *×* 10*^−^*^2^). Our results suggest that stabilizing selection is a pervasive force shaping the human plasma proteome and demonstrate that integrating genetic architecture with fitness-related consequences can connect long-term evolutionary constraint with contemporary phenotypic variation.

## 1 Introduction

Evolutionary processes continuously shape patterns of genetic and phenotypic variation within populations. While mutation introduces new genetic variants^1,2^, other forces such as drift, migration, and non-random mating influence their distribution over time^3,4,5,6,7^. Among these forces, natural selection impacts genetic variation through the association between genetic variants and fitness. Its signatures, embedded in both allele frequencies and phenotypic variation, provide insights into the evolutionary forces shaping human biology. Depending on how a phenotype influences fitness, selection may take different forms^8^. Directional selection shifts the population’s phenotypic mean towards a new optimum^9^, while disruptive selection favors extreme phenotypic values over intermediate ones^10^. In contrast, stabilizing selection, which selects against phenotypic extremes and maintains the population mean near an optimum, has long been proposed to be a predominant mode of selection shaping quantitative traits. Classic examples, such as human birthweight, for which both very low and very high values are associated with reduced neonatal survival^11^, illustrate how deviations from an optimum can be strongly deleterious. Beyond these complex phenotypes, evidence suggests that stabilizing selection is pervasive across different biological scales, acting on complex traits themselves and on the molecular traits that underlie them^12,13^.

Among molecular traits, proteins are the primary effectors of cellular function, acting as enzymes, receptors, and signaling molecules^14^. Their expression levels are expected to evolve around fitness optima because both under- and over-expression can be deleterious^13^. Consistent with this, genes encoding dosage-sensitive proteins are subject to strong evolutionary constraint, reflecting the fitness costs of perturbing optimal abundance levels^15^. In addition, highly abundant proteins tend to evolve more slowly at the sequence level, indicating strong functional constraint^16^. Protein abundance may therefore provide a biologically interpretable readout of evolutionary constraint. Changes in protein abundance can directly alter the stoichiometry of molecular complexes, the flux through metabolic and signaling pathways, and the balance between competing biochemical processes. Indeed, circulating protein levels have been linked to metabolic disease, neurodegeneration, cardiovascular risk, and immune dysfunction^17,18^, placing them at the intersection of molecular biology, physiology, and human health. Protein dosage sensitivity is also directly relevant to drug development, as therapeutically perturbing proteins whose levels are under strong constraint may carry a greater risk of adverse effects^19^.

Despite this central role, evolutionary analyses have historically focused on DNA sequence variation and the evolution of complex traits. Molecular intermediates have received less attention^13,20^, and existing work has largely addressed gene expression^21,22^ or metabolite levels^23^ rather than protein levels. To date, two broad classes of approaches have been developed to detect and quantify selection acting on quantitative traits. Microevolutionary methods investigate selection within populations over short timescales by testing trait-fitness correlations^24,25^, experimentally manipulating traits to observe fitness consequences^26^, tracking trait distributions across generations^27,28,29^, or detecting DNA-level signatures of selection within species^30^. Macroevolutionary approaches instead leverage divergence across species over longer timescales. These rely on comparative genomic analyses^31,32^ and phylogenetic modeling^33,34,35^ to identify signatures of long-term evolutionary constraints.

Under stabilizing selection, genetic variance persists through mutation-selection balance, and the relationship between effect size and allele frequency can be exploited to infer the strength of selection^12,36,37,38,39,40^. While all these approaches have advanced our understanding of adaptation and evolutionary mechanisms, they capture different modes of selection on different timescales, rely on distinct modeling assumptions and datasets, and their conclusions are rarely compared directly^12,41^. Bridging these approaches at the molecular level represents both a methodological and biological opportunity to gain deeper insights. Recent advances in large-scale plasma proteomics have made this feasible. The UK Biobank Pharma Proteomics Project (UKB-PPP) has measured thousands of circulating proteins in tens of thousands of individuals^42^, providing unprecedented statistical power to study evolutionary constraint on molecular phenotypes.

Here, we leverage the UKB-PPP resource to triangulate evidence of stabilizing selection^12,13,20,41^ acting on human plasma protein levels using two complementary approaches. This extends previous analyses of selection on molecular traits to the plasma proteome, and allows genetic-architecture and trait-fitness estimates of selection to be compared on the same traits. A graphical overview of the study design, datasets, and analytical framework is shown in Figure 1. First, we exploit the signature that stabilizing selection may leave on genetic architecture: variants with larger phenotypic effects are expected to be purged more efficiently, generating a negative relationship between allelic effect size and minor allele frequency (MAF). Second, we estimate selection from trait-fitness relationships, testing for an inverted U-shaped relationship between protein levels and fitness-related proxies capturing different dimensions of reproductive success, health, and longevity^43,25^. Finally, we compare our selection estimates with independent metrics of evolutionary constraint at the gene level. Within this framework, this work aims to address: (1) which human plasma proteins exhibit robust signatures of stabilizing selection, and (2) to what extent different methodological approaches produce concordant estimates of selection acting on proteins. These analyses provide an evolutionarily grounded framework for understanding protein dosage sensitivity, with broader implications for the molecular basis of human disease and the selection of drug targets.

**Figure 1:**
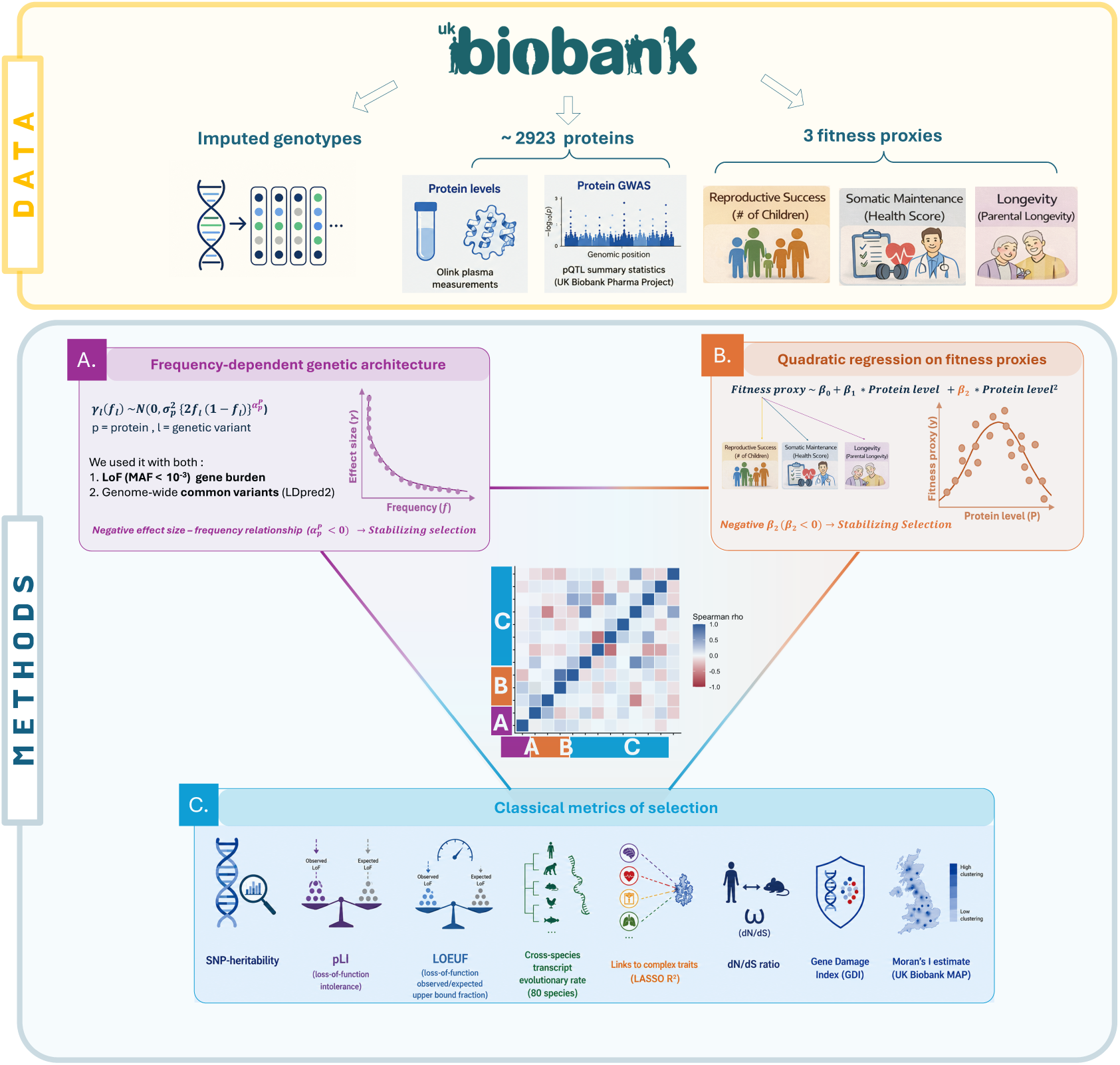
Graphical overview of the study design for inferring stabilizing selection on plasma protein levels. All data were derived from the UK Biobank. Inputs comprised imputed genotypes; plasma levels of 2’923 proteins from the UK Biobank Pharma Proteomics Project, used both as measured levels and as protein GWAS summary statistics; and three fitness proxies capturing complementary dimensions of fitness: reproductive success (number of children), somatic maintenance (composite health score), and lifespan potential (parental longevity). Two complementary frameworks were used to detect signatures of stabilizing selection on protein abundance. **(A)** Genetic architecture models quantified the relationship between allele frequency and effect size, using rare loss-of-function burden analyses and genome-wide common-variant models (LDpred2-auto). Under stabilizing selection, variants with larger effects are expected to remain at lower frequencies, yielding negative selection coefficients. **(B)**Quadratic regression models assessed the relationship between protein abundance and each fitness proxy; a significantly negative quadratic coefficient indicates an inverted U-shaped fitness landscape consistent with stabilizing selection. **(C)** Selection estimates were compared across approaches and with independent evolutionary and functional constraint metrics to assess their concordance and biological relevance.

## 2 Materials & Methods

### 2.1 Data pre-processing

#### Genotype and imputation

Individual genotype data were obtained from the UK Biobank^44^. Standard quality control procedures were applied as described by Bycroft et al.^45^, including filtering based on call rate, minor allele frequency, Hardy-Weinberg equilibrium, and imputation quality. Genotypes were imputed using the Haplotype Reference Consortium and UK10K/1000 Genomes reference panels^46^. For downstream polygenic modeling (e.g., LDpred2), we restricted analyses to high-quality HapMap3+ variants and used a European-ancestry LD reference panel consistent with the study sample^38^.

#### Whole-genome sequencing and gene burden scores

Whole-genome sequencing (WGS) data from the UK Biobank^47^ were used to construct rare variant gene burden scores. High-confidence loss-of-function (LoF) variants were aggregated at the gene level to generate burden instruments reflecting genetically proxied gene inactivation. Based on prior optimization analyses, a minor allele frequency (MAF) threshold of 1 *×* 10*^−^*^3^ was selected for downstream analysis. Additional methodological details are provided in section 2.2.2.

#### Sample selection

We applied standard sample quality control filters to the full UK Biobank genotyped cohort (488’377 individuals). Related individuals, defined by a kinship coefficient greater than 0.0442 in the UK Biobank kinship file, were excluded while retaining one representative per related group (407’219 individuals left). To reduce potential confounding from population stratification, we then restricted all analyses to participants with concordant self-reported and genetically inferred White British ancestry (337’545 individuals left). Records associated with redacted identifiers or with participants who had withdrawn consent before the start of our analyses (21 August 2023) were removed, resulting in an analytic cohort of 337’386 individuals.

#### Proteomics data

We analyzed Olink proteomics measurements for 53’061 UK Biobank participants across 2’923 proteins^42^. All analyses were restricted to the baseline visit (instance index = 0) and intersected with the previously defined QC-filtered, unrelated White British subset, resulting in n=36’185 samples with proteomics data. For each protein, we also used genome-wide association study (GWAS) summary statistics from the UK Biobank Pharma Proteomics Project (Olink Explore panels). Summary data were harmonized to a HapMap3+ reference panel (allele alignment and strand checks) and restricted to autosomal variants with minor allele frequency *>* 0.01. We used precomputed European-ancestry LD matrices^38^ for downstream modeling and, where required, aligned summary statistics to the LD reference panel (including sign flips for mismatched alleles). These harmonized pQTL summary statistics provided the inputs for downstream LDpred2-auto analyses of allele frequency-dependent genetic architecture for each protein.

#### Complex trait data

We curated a panel of 48 complex traits from UK Biobank phenotyping data (Supplementary Figure S1). The included traits were: body mass index (BMI), basal metabolic rate (BMR), body fat mass, height, hip circumference, waist circumference, weight, diastolic blood pressure, heart rate, systolic blood pressure, fluid intelligence, neuroticism, insulin-like growth factor 1 (IGF-1), sex hormone-binding globulin (SHBG), vitamin D, glycated hemoglobin (HbA1c), glucose, mean corpuscular hemoglobin (MCH), red blood cell (RBC) count, white blood cell (WBC) count, lymphocyte count, monocyte count, neutrophil count, platelet count, reticulocyte count, apolipoprotein A (ApoA), apolipoprotein B (ApoB), high-density lipoprotein cholesterol (HDL), low-density lipoprotein cholesterol (LDL), lipoprotein(a), triglycerides (TG), total cholesterol, alkaline phosphatase (ALP), alanine aminotransferase (ALT), aspartate aminotransferase (AST), gamma-glutamyltransferase (GGT), bilirubin, albumin, calcium (Ca), phosphate, C-reactive protein (CRP), total protein, forced vital capacity (FVC), creatinine (Cr), cystatin C, urea, urate, and right-hand grip strength. For each field, we retrieved all assessment instances, recoded non-informative responses (“do not know”, “prefer not to answer”) and negative codes as missing, and averaged across instances to obtain one value per participant. Educational attainment (EA) was also considered alongside the other complex traits. It was constructed from the UK Biobank qualification field (field 6138) by mapping categories to ISCED-equivalent years:

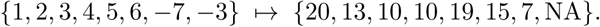

We then binarized EA as:

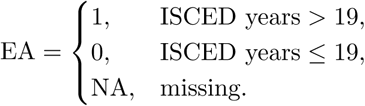

Traits were then merged with a covariate set (age, sex, genotyping batch, and the first 40 genetic principal components). The resulting dataset contained harmonized phenotypes spanning the anthropometric, cardiometabolic, hematological, renal, hepatic, and cognitive domains (Supplementary Figure S1).

#### Fitness proxies

We used three fitness proxies, each targeting a particular aspect of the reproductive cycle:

1. Number of children (direct reproductive success): we derived individual reproductive success proxies by extracting UK Biobank fields 2405 (number of children fathered) and 2734 (number of live births), recoding “do not know” and “prefer not to answer” responses as missing, and averaging across all available assessment instances to yield a single count per participant. This proxy captures realized reproductive output. Under classic Darwinian selection, higher offspring counts translate into greater fitness.
2. Composite health score (somatic maintenance): we took the 48 QC-filtered complex traits and adjusted each for the aforementioned covariates to obtain residual values that were used in the following analyses. To minimize redundancy, we pruned the 48 residual traits based on pairwise Spearman correlations, removing traits with an absolute correlation exceeding 0.4. This procedure retained 25 semi-independent traits: albumin, ALP, ALT, bilirubin, ApoA, ApoB, lipoprotein(a), glucose, HbA1c, Cr, urea, urate, systolic blood pressure, heart rate, BMI, phosphate, vitamin D, IGF-1, RBC count, WBC count, platelet count, reticulocyte count, right-hand grip strength, fluid intelligence, and neuroticism. The retained trait residuals were then scaled (mean = 0, SD = 1), squared and averaged across available traits, yielding a composite health score for each participant. The score captures an individual’s aggregate deviation from the population optimum across all traits, with low values representing high fitness. To facilitate interpretation and maintain consistency across fitness proxies, we inverted the final score so that higher values indicate smaller deviations from population-average phenotypes, potentially reflecting better somatic maintenance. This proxy relies on the assumption that the contemporary population mean approximates the evolutionary optimum for many traits. While this may not hold for each individual trait, we reasoned that, because the population has been continuously shaped by natural selection, the population mean should, across a broad range of traits, retain information about their evolutionary optima. Generally, more robust somatic health not only prolongs reproductive span and personal survival, but also underpins the immunity and resilience needed to raise and protect offspring through critical life stages, hence improving offspring fitness.
3. Parental longevity (lifespan potential): we derived a parental longevity phenotype using UK Biobank fields for age at death of both parents (father: field 1807; mother: field 3526). We recoded “do not know” and “prefer not to answer” responses as missing, inverse-normal-quantile transformed each age value before averaging maternal and paternal scores, yielding a single parental longevity score. Beyond one’s own childbearing, living into older age can improve the survival and reproductive success of grandchildren and other kin. By using parental longevity as a proxy for lifespan potential, we tap into the capacity to provide resources, knowledge and care long after reproduction has ceased, thereby enhancing the fitness of subsequent generations.

### 2.2 Detecting stabilizing selection on protein abundance

Stabilizing selection maintains quantitative traits near an optimal value by removing variants that cause large deviations, leaving two key signatures. First, at the genetic level, large-effect variants are constrained to low frequency, producing a negative relationship between allele frequency and squared effect size. Second, at the phenotypic level, both low and high trait values are associated with reduced fitness, generating an inverted U-shaped relationship between trait value and fitness. To detect stabilizing selection on plasma protein levels, we leveraged both signatures using two complementary approaches: (i) frequency-dependent genetic architecture models, including a rare-variant gene-burden framework and the genome-wide LDpred2-auto model for common variants, and (ii) quadratic regression of fitness proxies (reproductive success, somatic maintenance, and longevity) on observed protein levels.

#### 2.2.1 Frequency-dependent variant effects: genome-wide common variants

As outlined above, under stabilizing selection, variants with larger phenotypic effects are expected to be maintained at lower allele frequencies because extreme phenotypes are disadvantageous. This generates a negative relationship between effect size and allele frequency, which can be exploited to infer signatures of selection from genetic architecture. To capture this signal, we applied LDpred2-auto^38^, a method originally developed and extensively applied in the context of complex traits. To our knowledge, however, it has not previously been systematically applied to molecular traits such as protein abundance. For a given protein, LDpred2-auto models the per-variant (multivariable) causal effects as:

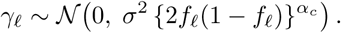

Here *f_ℓ_* is the allele frequency of variant *ℓ*, *γ_ℓ_* is the per-allele causal effect size of variant *ℓ* on the protein level, *σ*^2^ is a protein-specific parameter proportional to its narrow-sense heritability, and *α_c_* represents the selectionstrength coefficient for common variants. Values *α_c_ <* 0 indicate that variants with larger effects occur at lower frequency, consistent with stabilizing selection. To quantify *α_c_*, we applied LDpred2-auto^38^ to harmonized pQTL summary statistics from the UK Biobank Pharma Proteomics Project (Olink Explore panels). Analyses were restricted to harmonized HapMap3+ variants with minor allele frequency *>* 0.01, and precomputed European-ancestry LD matrices were used throughout. To focus on genome-wide polygenic regulation rather than local gene-specific effects, we restricted analyses to trans-acting variants, excluding all variants within *±*1 Mb of the gene encoding each protein (cis-pQTLs). Cis variants often exhibit disproportionately large effects driven by local regulatory architecture and substantially differ from classical complex trait architectures^22^, potentially obscuring broader signatures of stabilizing selection acting on protein regulation. As a sensitivity analysis, we repeated the analysis including cis variants. LDpred2-auto accounts for linkage disequilibrium via an ancestry-matched reference and jointly estimates SNP-heritability, the fraction of causal variants, and the selection coefficient *α_c_*. We ran the model using 50 independent chains with the causal fraction initialized on a log grid from 10*^−^*^8^ to 0.2 (500 burn-in and 500 sampling iterations; *α_c_* constrained to [*−*5, 1]; LD shrinkage 0.95, sign jumps disallowed). Following Privé et al.^38^, we retained well-converged chains and report the median and 95% interval of *α_c_*, SNP-heritability, and the fraction of causal variants. This LDpred2-based estimate provides a first selection estimate for each protein.

#### 2.2.2 Frequency-dependent variant effects: loss-of-function rare-variant burden

While the LDpred2-based approach targets common variants, rare variants can also be leveraged to infer signatures of selection. To capture gene-perturbation effects, we focused on rare (predicted) loss-of-function (LoF) variants (with minor allele frequency, MAF *<* 1 *×* 10*^−^*^3^). LoF alleles were defined as those classified as *high-confidence LoF* (HC LoF) by LOFTEE^48^. For each gene, burden scores were constructed by aggregating LoF alleles in the UKB’s WGS data into a gene-level dosage score coded as 0, 1, or 2, based on the maximum number of rare LoF allele copies carried by each individual at a given position within the gene (see the max parameter of regenie’s *build-mask* option for details). Genes without any LoF variant, or for which the LoF variant was not available in the whole-genome sequencing data, were excluded from downstream analysis. Gene-based burden association analyses were then performed against protein levels using regenie v.4.1^49^. REGENIE step 1 was performed with UK10K genotypes, using only individuals and SNPs that passed the filtering recommended in REGENIE’s tutorial for UK Biobank analyses, i.e. minor allele frequency (MAF) *≥* 0.01, minor allele count (MAC) *≥* 100, genotype missingness *≤* 10%, Hardy–Weinberg equilibrium *P >* 1 *×* 10*^−^*^15^, and individual missingness *≤* 10%. REGENIE step 2 was performed using the WGS data, conditioning effect estimates on common pQTLs reported in Hawkes et al. 2025^50^ to ensure independence from common variants. REGENIE yielded gene-specific burden effect-size estimates 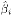, burden score frequencies *q_bi_*, and corresponding standard errors *se_bi_*.

Approximately 18’000 gene-level burden estimates were analyzed for each of the 2’923 proteins. We assumed that the observed burden effect estimates follow:

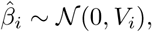

with variance:

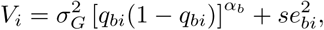

where 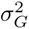 scales the overall genetic variance explained by rare variants, *q_bi_* denotes the burden allele frequency for gene *i*, and *α_b_* captures the relationship between effect size and allele frequency. The term 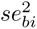 accounts for sampling variance from the burden association analysis. For each protein independently, parameters *α_b_* and 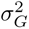 were estimated by maximizing the log-likelihood function:

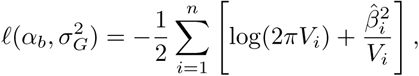

subject to the constraint:

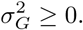

Optimization was performed independently for each protein, resulting in protein-specific estimates of frequency-dependent genetic architecture across the plasma proteome. Within this framework, the parameter *α_b_* provides an interpretable measure of selection acting on protein abundance. Negative values of *α_b_* indicate that larger burden effects tend to occur at lower frequencies, consistent with stabilizing selection removing extreme molecular phenotypes from the population. Values near *α_b_* = 0 indicate frequency-independent effect sizes consistent with neutral architectures, whereas positive *α_b_* values indicate larger effects at higher frequencies, a pattern theoretically compatible with either disruptive or balancing selection. This burden-based likelihood estimate provides a second selection estimate for each protein.

#### 2.2.3 Preprocessing of fitness proxies (covariate residualization)

For each fitness proxy *j ∈ {*n.children, health score, parental longevity*}*, we removed the variation explained by standard covariates. To achieve this, we regressed out covariates (age, age^2^, sex, genotyping batch, and genetic principal components PC1–PC40) and used the residuals in downstream analyses. For the number of children (n.children), we added two more covariates: educational attainment (EA) and the interaction EA*×*sex, because educational attainment is emerging as a strong determinant of an individual’s reproductive outcomes, especially for women (hence the interaction term)^51,52^.

#### 2.2.4 Quadratic regression of residualized fitness proxies on protein levels

Let *P_p_* be the vector of standardized baseline levels of protein *p* (*p* = 1, …, 2923) in our study sample. For each fitness proxy *Z_j_* (where *j* can represent n.children, health score, or parental longevity) and protein *p* we fit:

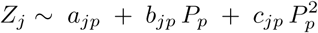

where *a_jp_* is the intercept term and *b_jp_, c_jp_* are the linear and quadratic coefficients. Under stabilizing selection on protein abundance (with all fitness proxies coded so that larger values indicate higher fitness), we expect an inverted U-shaped relationship (approximated by a downward-opening parabola), i.e., *c_jp_ <* 0, with optimal protein level:

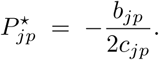

A significantly negative *c_jp_* indicates that both extremely low and extremely high protein levels are associated with worse fitness proxy values, consistent with stabilizing selection. Thus, for each protein *p* and fitness proxy *j* (n.children, health score, or parental longevity), the quadratic coefficient *c_jp_* serves as the selection estimate. A similar quadratic regression framework was recently applied by Souaiaia et al.^53^ to detect signatures of natural selection on different complex traits through associations with lifetime reproductive success.

### 2.3 Concordance between inferred selection estimates and independent evolutionary constraint metrics

To assess whether our five inferred selection coefficients capture meaningful evolutionary constraint, we sought to compare them against a series of complementary evolutionary and functional metrics using Spearman’s rank correlation. Proteins inferred to be under stronger stabilizing selection should show greater constraint across established measures of evolutionary and functional constraint. We considered nine different measures, capturing distinct dimensions of constraint:

1. **SNP-heritability:** Stabilizing selection favors intermediate phenotypes by selecting against deviations from an optimal trait value. Over evolutionary timescales, this process constrains additive genetic variation. Although new mutations continuously introduce new genetic variation, the amount of standing additive genetic variance is expected to reach a mutation-selection balance, with traits under stronger stabilizing selection exhibiting lower levels of additive genetic variance^39,54,55^. Consequently, proteins under stronger stabilizing selection are expected to have lower SNP-heritability (*h*^2^), as common genetic variants account for a smaller proportion of variation in their circulating levels. We quantified SNP-heritability for each protein using LDpred2-auto.
2. **pLI (loss-of-function intolerance)**^56^: it estimates the probability that a gene is intolerant to heterozygous loss-of-function (LoF) variation. A higher pLI value indicates that LoF variants in a given gene are strongly selected against, reflecting a greater degree of functional constraint. Consequently, proteins encoded by LoF-intolerant genes are expected to exhibit stronger signatures of stabilizing selection, as disruptive mutations are less likely to be tolerated in the population. pLI values were retrieved from the gnomAD database^56^.
3. **LOEUF**^56^: it is a continuous metric that quantifies the degree of constraint against loss-of-function (LoF) variants. It represents the upper bound of the ratio between observed and expected LoF variants for a gene, with lower values indicating a greater depletion of such variants. Genes with low LOEUF scores are considered LoF-intolerant. As a result, proteins encoded by these highly constrained genes are expected to exhibit stronger signatures of stabilizing selection. LOEUF values were retrieved from the gnomAD database.
4. **Cross-species transcript evolutionary rate**: transcript evolutionary rates quantify divergence of gene expression levels over the course of evolution. To estimate the evolutionary conservation of mammalian gene expression, we used RNA-seq data from Liu et al.^57^, comprising transcript levels of 13’452 protein-coding genes measured in brain, kidney, and liver across 103 mammalian species. Analyses were restricted to genes that met three criteria: (1) the encoded proteins were included in the UK Biobank Olink panel, (2) the genes were measured in at least 70% of all samples, and (3) they were measured in at least 80 species. Organ-specific evolutionary rates were estimated for 1’160, 1’142, and 1’085 proteins in brain, kidney, and liver, respectively. Gene expression values were reported as log_2_-transformed RPKM values normalized using the trimmed mean of M-values method, as described previously^57^. Evolutionary rates were calculated by fitting a Brownian Motion (BM) model of trait evolution to transcript levels for each gene and organ across the phylogeny, using the fitContinuous function in the *geiger* R package^58^. The BM model treats trait evolution as a random walk along phylogenetic branches, where, over an elapsed time Δ*t*, the trait value *x* changes by a random value drawn from a normal distribution with mean 0 and variance *σ*^2^Δ*t*:

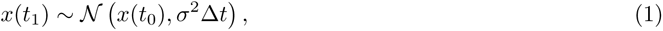

where *x*(*t*_0_) is the ancestral trait value, *x*(*t*_1_) is the expected trait value after time Δ*t*, and *σ*^2^ is the evolutionary rate describing how quickly trait variance accumulates over time. Using the known phylogenetic tree and the observed trait values at the tips, we estimated *σ*^2^ following Felsenstein’s method^59^. This allowed us to quantify the rate of trait diversification while accounting for shared evolutionary history among species. The evolutionary rate, *σ*^2^, is expressed as the increase in trait variance per unit of evolutionary time, approximated by phylogenetic branch length, with lower values potentially indicating stronger long-term stabilizing selection acting on gene expression across species. In our analyses, organ-specific rates were standardized within each tissue and combined into a single summary measure using inverse-variance weighted meta-analysis. Lower evolutionary rates reflect stronger long-term stabilizing selection acting on gene expression across species. Importantly, evolutionary constraint observed across species is expected to mirror constraint within species: genes whose expression levels are tightly conserved over long evolutionary timescales are likely to tolerate only limited variation in expression within populations. According to the neutral theory of molecular evolution, the same functional requirements that restrict divergence between species also constrain variation within species^60^. Consequently, genes with highly conserved expression profiles are expected to be under stronger selective constraint at multiple levels^23^. In turn, the proteins encoded by these genes are predicted to exhibit stronger signatures of stabilizing selection.
5. **Links to complex traits (LASSO** *R*^2^): it quantifies the extent to which variation in protein abundance can be explained by a panel of complex traits, spanning multiple biological domains. For each protein, we fitted a LASSO regression of protein levels on the 48 standardized complex traits described above, selecting the penalty by 10-fold cross-validation (cv.glmnet, *λ*_min_). The cross-validated *R*^2^ was used as a measure of explanatory power, with higher values indicating stronger coupling between protein levels and complex phenotypes. Under the hypothesis that fitness acts on higher-level phenotypes, complex traits represent a primary target of selection. Complex traits are themselves subject to stabilizing selection^36^, whereby deviations from an optimal range are selected against. Consequently, proteins whose abundance is more strongly associated with complex traits under stabilizing selection are expected to experience stronger indirect selective constraint.
6. **dN/dS ratio** represents the ratio of non-synonymous to synonymous substitution rates (dN/dS) which measures coding-sequence evolutionary constraint across species. Lower dN/dS values potentially indicate stronger purifying selection and greater protein-coding sequence conservation, whereas higher values suggest weaker constraint or accelerated sequence evolution. We computed dN/dS values for human-mouse orthologous genes using the orthologr R package^61^, which provides a framework for genome-wide orthology inference and dN/dS estimation. Orthologous genes were identified between human and mouse, and synonymous and non-synonymous substitution rates were estimated for each orthologous gene pair. Proteins encoded by genes with lower dN/dS values are expected to show stronger stabilizing selection signatures, reflecting stronger long-term evolutionary constraint on protein-coding sequence.
7. *ω_A_*: it is a measure of adaptive evolution at the gene level, using the estimates provided by Latrille et al. (2023) for mammalian protein-coding genes^62^. In contrast to the conventional dN/dS, which measures the overall rate of non-synonymous substitution relative to synonymous substitution, *ω_A_* = dN/dS *− ω*_0_ is designed to quantify the adaptive component of sequence evolution by subtracting the expectation under a nearly neutral model (*ω*_0_). This explicit nearly neutral model is rooted in mutation-selection models, which represent site-specific amino-acid fitness differences and therefore provide a more mechanistic description of purifying selection than a neutral dN/dS = 1 reference^63,64,65^. The inclusion of *ω_A_* is therefore complementary to dN/dS: whereas dN/dS summarizes total non-synonymous evolutionary constraint, *ω_A_* allows for identifying genes experiencing positive selection despite overall purifying selection^64,62^. Although elevated *ω_A_* reflects rapid amino-acid turnover across mammals, its relationship with stabilizing selection on human protein abundance remains unknown.
8. **Gene Damage Index (GDI)**: it quantifies the cumulative mutational damage accumulated within human protein-coding genes in the general population and therefore serves as a gene-level measure of mutational tolerance and long-term functional constraint. Specifically, GDI summarizes the burden of non-synonymous variation observed across population sequencing datasets, with highly damaged genes interpreted as more tolerant to functional variation and therefore under weaker selective constraint. Conversely, genes with low GDI values harbor fewer accumulated damaging variants and are considered more functionally indispensable. As a result, proteins encoded by low-GDI genes are expected to exhibit stronger signatures of stabilizing selection acting on protein abundance. GDI estimates were retrieved from Itan et al. (2015)^66^.
9. **Moran’s *I* estimate**: Moran’s *I* was used to quantify the degree of spatial autocorrelation in protein abundance across study participants with known geographic location, measuring the extent to which similar protein levels cluster geographically. While Moran’s *I* is typically used to characterize spatial structure arising from demographic history, population structure, or assortative mating, it can also quantify geographic clustering of biological traits. Such clustering is expected to increase when variation in a trait is driven by geographically structured environmental factors^67^. To estimate Moran’s *I*, we first calculated pairwise Euclidean distances (*d_ij_*, in km) between study participants based on their geographic coordinates and transformed these distances into a spatial weights matrix using an exponential distance-decay function (*w_ij_* = exp 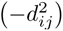). The resulting weights were then used to compute Moran’s *I* for each protein using the moran.test function from the R package *spdep*^68^.

## 3 Results

### 3.1 Classification of selection signatures inferred from protein-fitness relationships

Quadratic regression models (*p <* 0.05) enabled the classification of protein-fitness associations into five categories (Figure 2): (i) *directional-linear*, defined by a significant linear term and a non-significant quadratic term; (ii) *stabilizing*, characterized by a significant negative quadratic coefficient with the inferred optimum located within *±*1 SD of the protein distribution mean; (iii) *disruptive*, defined by a significant positive quadratic coefficient with the optimum within *±*1 SD; (iv) *directional non-linear (stabilizing-type)*, where the quadratic coefficient is negative but the inferred optimum lies outside the *±*1 SD range; and (v) *directional non-linear (disruptive-type)*, where the quadratic coefficient is positive and the inferred optimum falls outside the *±*1 SD range. Across all three fitness proxies, negative quadratic coefficients were substantially more frequent than positive coefficients, indicating an overall predominance of stabilizing-type curvature in protein-fitness relationships. This enrichment was particularly pronounced for the health score phenotype, for which a large proportion of proteins exhibited either classical stabilizing patterns or directional non-linear stabilizing-type relationships. In contrast, analyses based on parental longevity and number of children identified comparatively fewer proteins with centered optima and a greater proportion of directional-linear only associations. Importantly, only a subset of proteins exhibiting concave quadratic relationships satisfied the strict definition of classical stabilizing-type selection requiring an inferred optimum within *±*1 SD of the observed protein distribution mean. Many additional proteins instead exhibited displaced optima outside this central range, suggesting asymmetric or shifted fitness landscapes that may reflect ongoing convergence to a new population optimum. Overall, the marked excess of negative quadratic coefficients across all fitness proxies suggests that evolutionary pressures more commonly constrain protein abundance within relatively narrow functional ranges.

**Figure 2:**
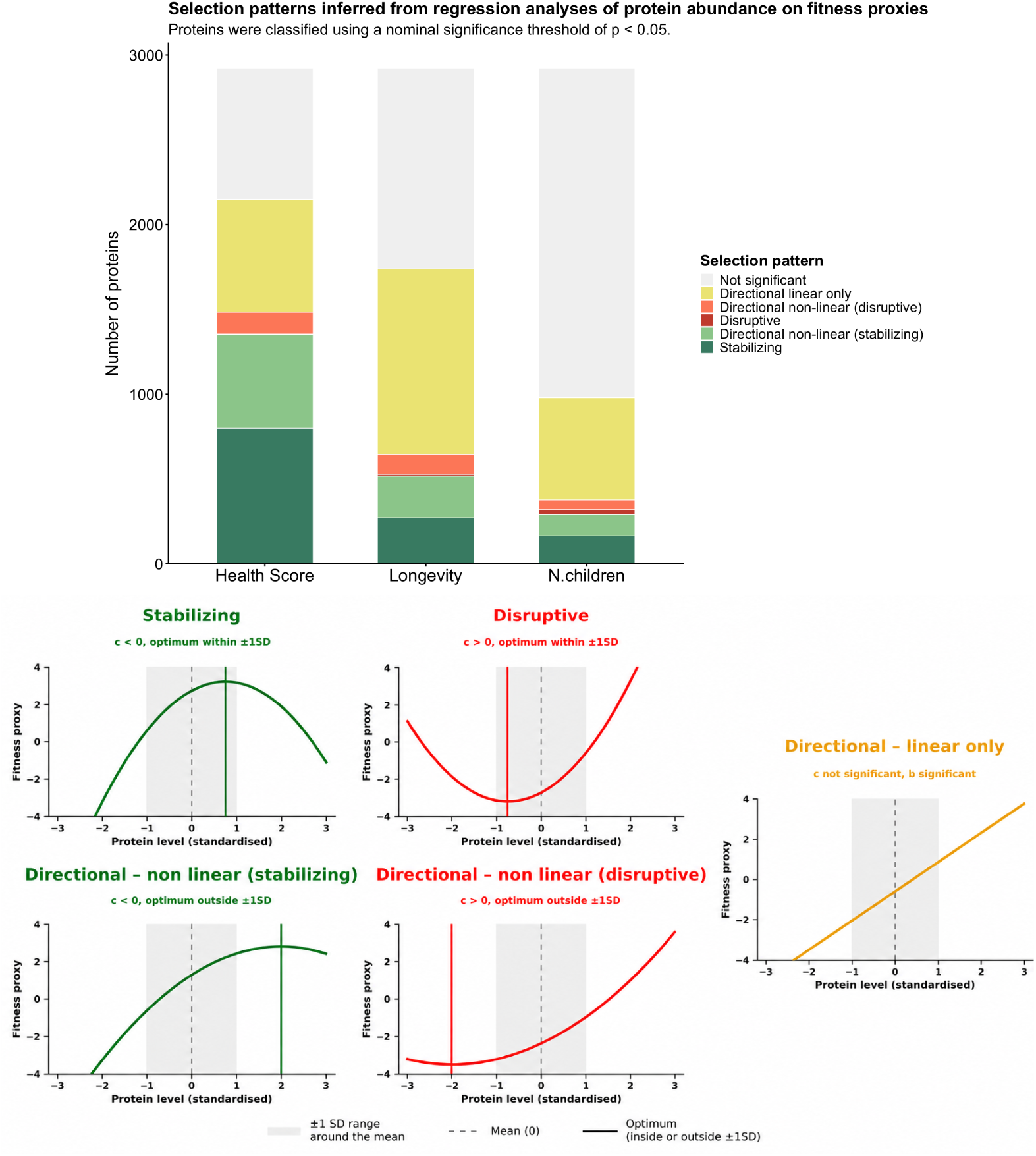
Classification of distinct signatures of selection acting on protein abundance inferred from quadratic regression models. **(A)** Counts of proteins assigned to selection categories from quadratic regressions of three fitness proxies (health score, parental longevity, and number of children) on standardized protein abundance. Stabilizing patterns are most frequent for the health score proxy, whereas directional associations dominate for longevity and number of children. However, among proteins with a significant quadratic term, stabilizing selection consistently represents a larger proportion than disruptive selection. **(B)** Schematic representation of each category showing the relationship between standardized protein levels and the fitness proxy. Categories are defined according to the significance and sign of the linear and quadratic terms: *stabilizing* (negative quadratic term with inferred optimum within *±*1 SD), *disruptive* (positive quadratic term with inferred optimum within *±*1 SD), *directional linear only* (significant linear term with non-significant quadratic term), *directional-non-linear* (significant quadratic term with optimum outside *±*1 SD), and *not significant*. Categories were defined using an in-range criterion of *±*1 SD for *P*\* and nominal significance (*p <* 0.05).

### 3.2 Multiple analytical frameworks reveal a predominance of stabilizing selection signatures across protein abundance traits

We summarized inferred selection coefficients across all analytical approaches, considering nominally significant estimates (*p <* 0.05) as a compromise between power and type I error. Because each inference framework relies on distinct assumptions and sensitivity profiles, we applied filters to retain only sufficiently reliable estimates. For LDpred2-derived coefficients, analyses were restricted to proteins with significantly non-zero estimates of heritability and polygenicity (*h*^2^ *>* 0.1, *p_causal_ >* 0.00025, *p_h_*2 *<* 0.05, *p_pcausal_ <* 0.05). For observational quadratic regression coefficients, analyses were restricted to proteins with inferred optima located within *±*1 standard deviation of the observed protein abundance distribution and with *h*^2^ *>* 0.1. The heritability filter ensures that inferred selection signals were estimated only for sufficiently heritable proteins. Indeed, selection acting on protein abundance cannot be reliably inferred from genetic data for traits with negligible heritability. As shown in Figure 3, negative selection coefficients substantially outnumbered positive coefficients across all analytical frameworks. Within the LDpred2 framework, 657 proteins displayed nominally significant negative frequency-dependence parameters (*α <* 0), indicating that 100% of significant associations were negative. A similar pattern was observed in the rare-variant burden analyses, in which 367 proteins exhibited significantly negative coefficients compared with only 30 proteins showing positive coefficients, corresponding to 92.4% negative associations. The predominance of negative coefficients became even more pronounced in the quadratic regression analyses after restricting to proteins with inferred optima within *±*1 SD of the observed distribution. For the health score phenotype, nearly all significant proteins (99.8%) exhibited negative quadratic coefficients consistent with stabilizing selection. Similarly, analyses based on parental longevity demonstrated strong enrichment for stabilizing patterns, with 97.7% of significant proteins displaying negative coefficients. Although the analysis based on number of children identified comparatively more disruptive-like signals than the others, stabilizing-like associations still represented the clear majority (89.3%). Together, these findings reveal a consistent predominance of negative coefficients across complementary analytical frameworks, supporting widespread signatures compatible with stabilizing selection on protein abundance. The same overall pattern remained consistent after multiple-testing correction using a false discovery rate (FDR) threshold, although the total number of significant proteins was reduced (Supplementary Figure S5). To assess robustness and specificity, we additionally performed sensitivity analyses for the number of children phenotype by excluding educational attainment as an additional covariate, and conducted negative-control analyses using tea intake, a phenotype not expected to directly reflect fitness. Importantly, this overall pattern remained highly consistent irrespective of whether educational attainment was included as a covariate for number of children (Supplementary Figure S3). In contrast, analyses using tea intake as a negative-control phenotype showed that only 47.2% of significant coefficients were negative (Supplementary Figure S3).

**Figure 3:**
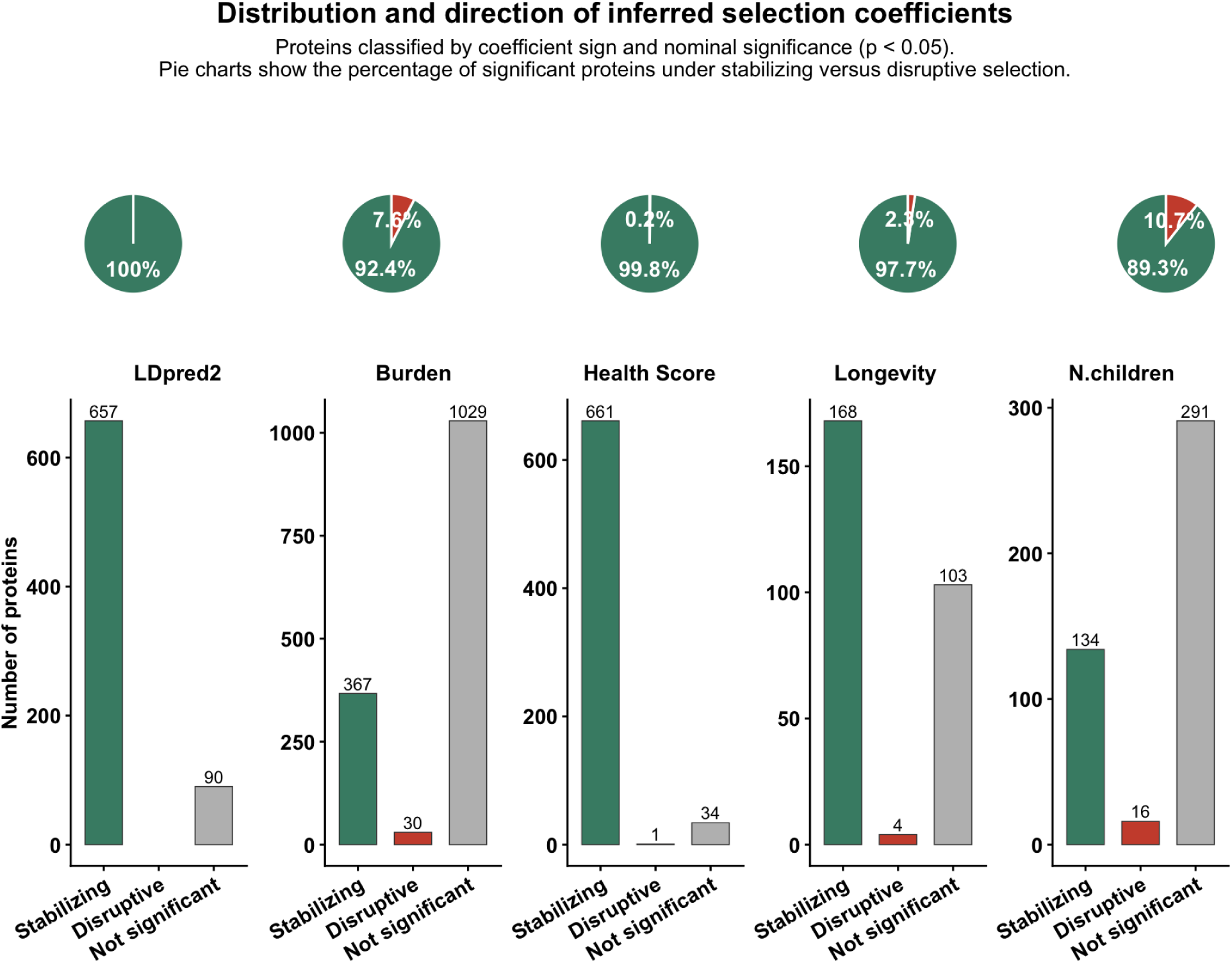
Distribution and direction of inferred selection coefficients across methods: clear predominance of stabilizing selection signatures across protein abundance traits. Proteins were classified according to the sign and significance of inferred selection coefficients into three categories: significant negative selection coefficients (*α_c_*, *α_b_*, or *c <* 0; nominal *p <* 0.05), significant positive coefficients (*α_c_*, *α_b_*, or *c >* 0), and non-significant proteins. Counts are shown for LDpred2-based estimates, rare variant burden analyses, and regression models relating observed protein abundance to three fitness proxies (health score, parental longevity, and number of children). Pie charts above each panel show the relative proportion of significant negative versus significant positive coefficients among proteins reaching nominal significance. For the regression analyses, only proteins passing the quality filters reported in the main text were retained. Across all analyses, significant negative coefficients substantially outnumbered significant positive coefficients, indicating a consistent predominance of stabilizing selection signatures.

### 3.3 Concordance between inferred selection coefficients across methodological frameworks

Next, we examined the concordance between selection coefficients inferred using the different methodological frameworks (Figure 4) to determine whether these approaches capture a shared underlying signal of stabilizing selection acting on protein abundance. Overall, correlations across methods were modest, suggesting that each framework captures partially distinct aspects of selective constraint. Nevertheless, several positive cross-method correlations emerged. The strongest concordance was observed between health score-derived quadratic coefficients and rare-variant burden estimates (nominally significant estimates: *ρ* = 0.36, *p* = 5.2 *×* 10*^−^*^6^). Health score-derived coefficients additionally showed a weaker positive correlation with LDpred2-derived estimates among proteins passing 5% FDR threshold (*ρ* = 0.11, *p* = 3.4 *×* 10*^−^*^2^). When considering all proteins passing the quality-control and heritability filters, but without imposing significance thresholds on the inferred selection coefficients (Supplementary Figure S6), the selection coefficients derived from the quadratic model using the three different fitness proxies exhibited modest positive pairwise correlations (*ρ* = 0.19–0.26, *p* = 10*^−^*^2^–10*^−^*^4^). The UpSet plot further revealed substantial heterogeneity in the overlap of stabilizing selection signals across methods (Figure 5). The largest pairwise overlap was observed between LDpred2- and health score-based analyses (220 proteins). LDpred2-, burden-, and health score-derived analyses uniquely identified 163, 159, and 149 proteins, while 62 proteins were shared across all three. Higher-order overlaps were progressively less frequent. Five proteins showed consistent stabilizing-like signatures across all five analyses: GPC1, ACAN, CDH3, IGFBP3 and SMPDL3A (Table 2). These proteins participate in extracellular matrix organization, growth-factor signaling, epithelial adhesion, and cellular homeostasis, processes linked to tissue integrity, maintenance, and developmental stability. Overall, these findings provide complementary, partially convergent evidence consistent with stabilizing selection acting on protein abundance. To investigate the biological functions associated with inferred selection signatures, we performed Gene Ontology (GO) enrichment analyses (Supplementary Figure S9). Proteins identified by the rare-variant burden analysis were enriched for lysosomal, vacuolar, and secretory-granule compartments, suggesting selective constraints on proteins involved in intracellular degradation and regulated secretion. Among the fitness-proxy analyses, parental longevity showed enrichment for cell–cell adhesion.

**Table 1:** Cross-method correlations between inferred selection coefficients under nominal, FDR-filtered, and unfiltered analyses.

| Variable 1 | Variable 2 | $\rho$ (Nominal) | $P$ (Nominal) | $\rho$ (FDR) | $P$ (FDR) | $\rho$ (No filter) | $P$ (No filter) | Significance |
| --- | --- | --- | --- | --- | --- | --- | --- | --- |
| Burden | Health Score | 0.365 | $5.16 \times 10^{-6}$ | 0.359 | 0.0045 | 0.246 | $5.47 \times 10^{-8}$ | Nominal & FDR & No filter |
| Burden | Longevity | -0.200 | 0.290 | — | — | 0.149 | 0.0447 | No filter only |
| Health Score | Longevity | 0.058 | 0.511 | 0.027 | 0.853 | 0.260 | $2.05 \times 10^{-4}$ | No filter only |
| Health Score | N.children | 0.031 | 0.756 | 0.046 | 0.818 | 0.235 | $1.14 \times 10^{-4}$ | No filter only |
| LDpred2 | Health Score | 0.076 | 0.127 | 0.107 | 0.0344 | -0.027 | 0.555 | FDR only |

**Figure 4:**
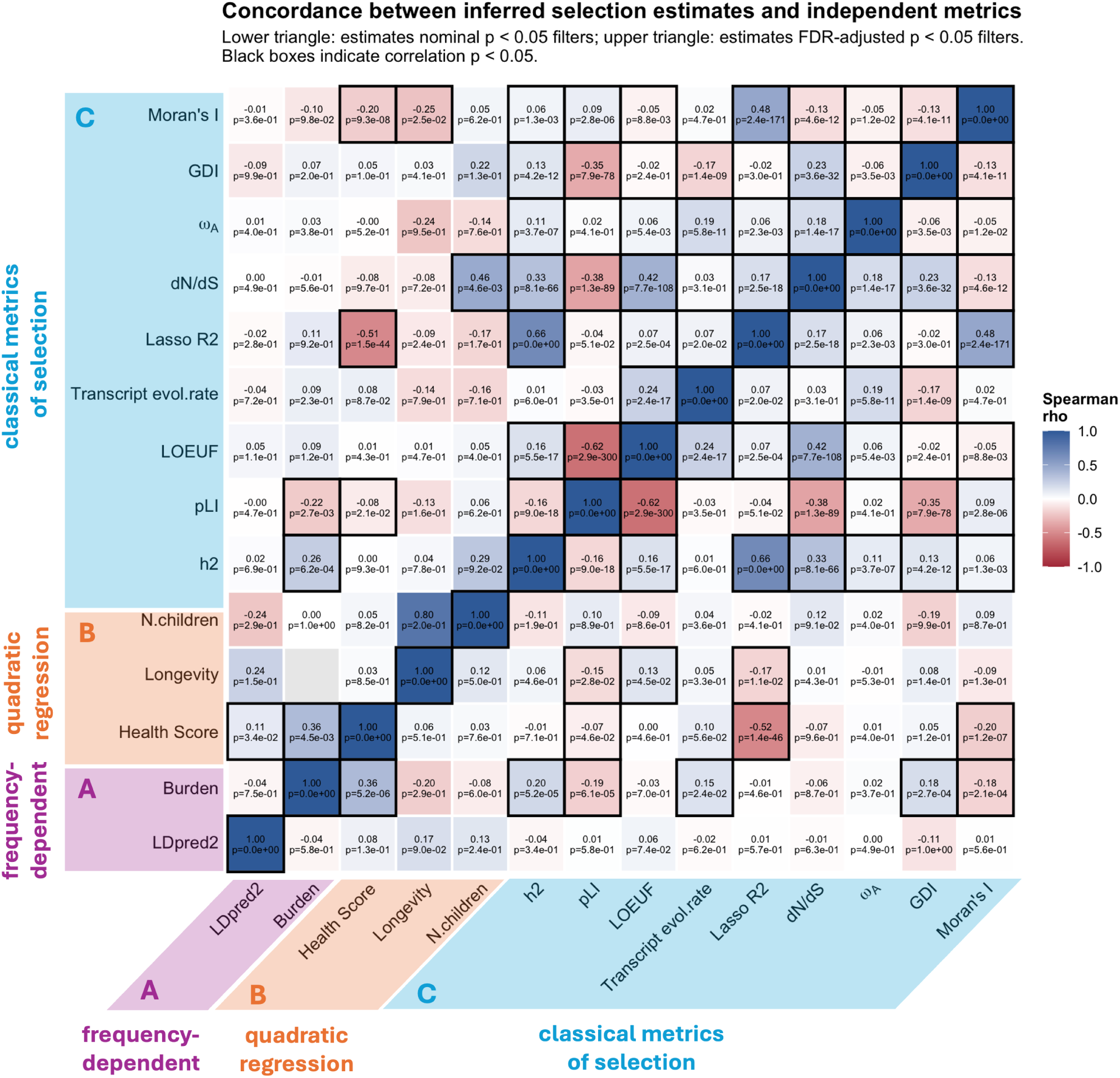
Concordance between inferred protein-level selection coefficients and independent measures of evolutionary and functional constraint. Heatmap showing Spearman correlations between inferred protein-level selection coefficients and independent measures of evolutionary and functional constraint. Following the triangulation framework summarized in the graphical abstract, analyses were organized into three complementary categories: (A, purple) frequency-dependent genetic architecture estimates derived from both common- and rare-variant approaches, (B, orange) regression-based selection estimates derived from quadratic models using fitness proxies, and (C, blue) classical evolutionary and functional constraint metrics. The latter include protein heritability (*h*^2^), probability of loss-of-function intolerance (pLI), LOEUF, meta-transcript evolutionary rate, LASSO *R*^2^, dN/dS, *ωA*, Gene Damage Index (GDI), and Moran’s I. The lower triangle shows correlations computed using inferred selection coefficients passing nominal significance thresholds (*p <* 0.05), whereas the upper triangle shows correlations based on coefficients remaining significant after false discovery rate (FDR) correction. Cell colors indicate Spearman correlation coefficients (*ρ*), with blue representing positive correlations and red representing negative correlations. Black borders indicate nominally significant correlations (*p <* 0.05).

**Figure 5:**
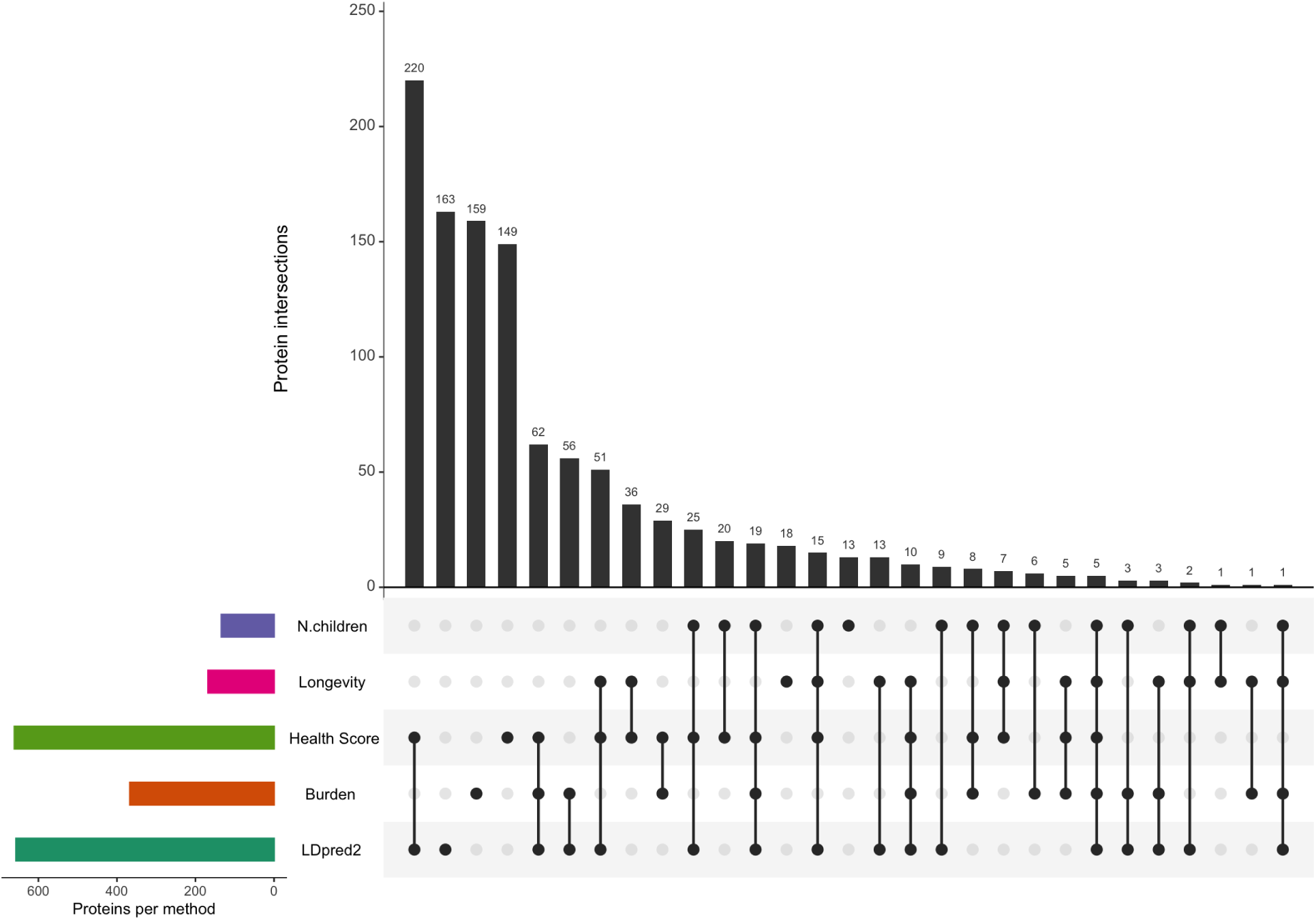
Overlap of proteins showing evidence of stabilizing selection across the different analytical frameworks. Vertical bars indicate the size of each protein intersection, while the connected dot matrix denotes the specific combination of methods supporting each overlap. Horizontal bars show the total number of proteins identified by each method. Several intersections showed substantial replication across analytical frameworks, including 220 proteins jointly identified by LDpred2 and Health Score analyses and 62 proteins overlapping between LDpred2, Burden, and Health-score. Five proteins were consistently supported across all five methods. Proteins were required to pass method-specific filtering criteria and reach nominal significance (*P <* 0.05).

**Table 2:** Proteins consistently supported across all five stabilizing-selection frameworks. These proteins, identified by all five methods, participate in biological processes related to tissue organization, growth-factor signaling, and cellular homeostasis.

| Protein | Functional role | Disease relevance | References |
| --- | --- | --- | --- |
| Glypican-1 | Heparan sulfate proteoglycan involved in growth-factor signaling, angiogenesis, and tissue morphogenesis. | Cancer progression, neurodevelopment, and vascular biology. | 69,70 |
| Aggrecan | Major cartilage extracellular matrix proteoglycan required for hydration and skeletal integrity. | Osteoarthritis, skeletal dysplasia, and growth disorders. | 71,72 |
| Cadherin-3 | Calcium-dependent adhesion protein important for epithelial organization and tissue architecture. | Hypotrichosis and aggressive epithelial cancers. | 73,74 |
| IGF-binding protein 3 | Major circulating carrier of IGF-I/II; regulates growth, apoptosis, metabolism, and aging. | Cancer susceptibility, metabolic dysfunction, and lifespan-related traits. | 75,76 |
| SMPDL3A | Membrane-associated phosphodiesterase involved in sphingolipid metabolism and cellular homeostasis. | Metabolic regulation and inflammatory signaling. | 77 |

### 3.4 Correlations between inferred selection estimates and independent constraint metrics

We next assessed the concordance between inferred protein-level selection coefficients and independent measures of evolutionary and functional constraint using Spearman correlations. The resulting correlation heatmap (Figure 4) summarizes concordance between inferred selection coefficients and independent evolutionary metrics after applying the filtering criteria described above. The lower triangle of the heatmap corresponds to analyses using only nominal significant inferred selection coefficients (*p <* 0.05), whereas the upper triangle corresponds to false discovery rate (FDR)-controlled thresholds. Black borders indicate nominally significant (*p <* 0.05) Spearman correlations. Supplementary Figure S7 additionally reports the number of overlapping proteins used for each pairwise correlation following the described filtering procedures. Overall, LDpred2-derived selection estimates showed relatively weak associations with the independent metrics considered here, and no robust correlations were consistently observed across different significance thresholds. In contrast, burden-based selection coefficients displayed substantially clearer and biologically coherent patterns: (i) they correlate positively with protein heritability (*h*^2^; nominal: *ρ* = 0.20, *p* = 5.2 *×* 10*^−^*^5^; FDR: *ρ* = 0.26, *p* = 6.2 *×* 10*^−^*^4^), indicating that proteins under stronger inferred stabilizing selection constraint are less heritable. (ii) They show negative correlations with pLI (nominal: *ρ* = *−*0.19, *p* = 6.1 *×* 10*^−^*^5^; FDR: *ρ* = *−*0.22, *p* = 2.7 *×* 10*^−^*^3^), suggesting that genes exhibiting greater intolerance to loss-of-function variation are under stronger stabilizing selection. (iii) Positive associations were also observed with transcript evolutionary rate (nominal: *ρ* = 0.15, *p* = 2.4 *×* 10*^−^*^2^), GDI (nominal: *ρ* = 0.18, *p* = 2.7 *×* 10*^−^*^4^), and Moran’s I (nominal: *ρ* = *−*0.18, *p* = 2.0 *×* 10*^−^*^4^). Selection inferred using quadratic models with health score and parental longevity as fitness proxies also showed partial agreement with independent constraint metrics.

Health score-derived estimates correlated with (i) LASSO *R*^2^ (nominal: *ρ* = *−*0.52, *p* = 1.4 *×* 10*^−^*^46^; FDR: *ρ* = *−*0.51, *p* = 1.5 *×* 10*^−^*^44^); (ii) pLI (nominal: *ρ* = *−*0.07, *p* = 4.6 *×* 10*^−^*^2^; FDR: *ρ* = *−*0.08, *p* = 2.1 *×* 10*^−^*^2^); and (iii) Moran’s I (nominal: *ρ* = *−*0.20, *p* = 1.2 *×* 10*^−^*^7^; FDR: *ρ* = *−*0.20, *p* = 9.3 *×* 10*^−^*^8^). Similarly, longevity-derived estimates showed associations with pLI (nominal: *ρ* = *−*0.15, *p* = 2.8 *×* 10*^−^*^2^), LOEUF (nominal: *ρ* = 0.13, *p* = 4.5 *×* 10*^−^*^2^), and Moran’s I (FDR: *ρ* = *−*0.25, *p* = 2.5 *×* 10*^−^*^2^). Number of children-derived estimates correlated positively with dN/dS (*ρ* = 0.46, *p* = 4.6 *×* 10*^−^*^3^). In addition, we repeated the full correlation analysis without applying any significance threshold to the inferred selection coefficients. The corresponding correlation heatmaps and overlap matrices are provided in Supplementary Figures S6 and S8.

**Table 3:** Correlations between metrics of selection and external independent constraint metrics under nominal, FDR, and unfiltered analyses.

| Variable 1 | Variable 2 | $\rho$ (Nominal) | $P$ (Nominal) | $\rho$ (FDR) | $P$ (FDR) | $\rho$ (No filter) | $P$ (No filter) | Significance |
| --- | --- | --- | --- | --- | --- | --- | --- | --- |
| Burden | GDI | 0.177 | $2.73 \times 10^{-4}$ | 0.066 | 0.202 | -0.045 | 0.949 | Nominal only |
| Burden | Lasso $R^2$ | -0.005 | 0.459 | 0.107 | 0.919 | -0.242 | $1.02 \times 10^{-20}$ | No filter only |
| Burden | Moran's $I$ | -0.177 | $2.06 \times 10^{-4}$ | -0.099 | 0.0977 | -0.115 | $6.62 \times 10^{-6}$ | Nominal & No filter |
| Burden | Transcript evol. rate | 0.151 | 0.0240 | 0.086 | 0.232 | 0.005 | 0.456 | Nominal only |
| Burden | $h^2$ | 0.202 | $5.23 \times 10^{-5}$ | 0.258 | $6.18 \times 10^{-4}$ | -0.191 | $4.21 \times 10^{-13}$ | Nominal & FDR & No filter |
| Burden | pLI | -0.194 | $6.10 \times 10^{-5}$ | -0.215 | 0.0027 | 0.043 | 0.944 | Nominal & FDR |
| Health Score | Lasso $R^2$ | -0.516 | $1.40 \times 10^{-46}$ | -0.510 | $1.51 \times 10^{-44}$ | -0.545 | $2.28 \times 10^{-55}$ | Nominal & FDR & No filter |
| Health Score | Moran's $I$ | -0.200 | $1.16 \times 10^{-7}$ | -0.203 | $9.26 \times 10^{-8}$ | -0.202 | $4.09 \times 10^{-8}$ | Nominal & FDR & No filter |
| Health Score | pLI | -0.067 | 0.0462 | -0.081 | 0.0208 | -0.070 | 0.0351 | Nominal & FDR & No filter |
| LDpred2 | LOEUF | 0.058 | 0.0736 | 0.050 | 0.106 | 0.071 | 0.0283 | No filter only |
| LDpred2 | $h^2$ | -0.037 | 0.339 | 0.016 | 0.686 | -0.239 | $3.89 \times 10^{-11}$ | No filter only |
| Longevity | LOEUF | 0.133 | 0.0454 | 0.010 | 0.471 | 0.254 | $1.69 \times 10^{-5}$ | Nominal & No filter |
| Longevity | Lasso $R^2$ | -0.174 | 0.0110 | -0.091 | 0.242 | -0.350 | $1.26 \times 10^{-9}$ | Nominal & No filter |
| Longevity | Moran's $I$ | -0.087 | 0.128 | -0.249 | 0.0255 | -0.123 | 0.0209 | FDR & No filter |
| Longevity | dN/dS | 0.014 | 0.433 | -0.077 | 0.716 | 0.169 | 0.00431 | No filter only |
| Longevity | pLI | -0.150 | 0.0282 | -0.132 | 0.160 | -0.165 | 0.00375 | Nominal & No filter |
| N.children | LOEUF | -0.091 | 0.860 | 0.047 | 0.399 | 0.107 | 0.0134 | No filter only |
| N.children | Lasso $R^2$ | -0.020 | 0.405 | -0.169 | 0.169 | -0.133 | 0.00263 | No filter only |
| N.children | dN/dS | 0.116 | 0.0909 | 0.460 | 0.00457 | 0.110 | 0.0148 | FDR & No filter |
| N.children | $h^2$ | -0.108 | 0.188 | 0.293 | 0.0924 | -0.168 | $4.03 \times 10^{-4}$ | No filter only |
| N.children | $\omega_A$ | 0.023 | 0.399 | -0.143 | 0.762 | 0.093 | 0.0414 | No filter only |

## 4 Discussion

By integrating genome-wide frequency-dependent genetic architecture (LDpred2), rare-variant burden modeling, and quadratic regression using multiple fitness proxies, we aimed to capture both long-term evolutionary constraint shaping genetic architecture and contemporary fitness consequences of protein level variation. The novelty of our study is threefold: (i) instead of complex traits, we focus on evolutionary mechanisms acting on molecular human readouts; (ii) we developed a rare-variant burden framework to estimate selection; and (iii) we compare a plethora of state-of-the-art approaches tackling orthogonal aspects of selection. A central finding of this study is the consistent predominance of signatures compatible with stabilizing selection across analytical approaches. Across both genetic and observational approaches, negative inferred selection coefficients substantially outnumbered positive coefficients, suggesting that extreme deviations from population mean levels are generally disfavored. This concordance across fundamentally distinct methodological frameworks provides convergent evidence that circulating protein levels are commonly maintained near functional optima. These results further support the broader hypothesis that stabilizing selection acts not only on complex phenotypes, but also extensively constrains molecular traits underlying physiological homeostasis. The link between the predictability of a protein using complex traits and the inferred strength of selection acting on it suggests that a large part of the evolutionary forces shaping protein abundance acts indirectly via complex phenotypes. The three fitness proxies we used capture distinct yet partially overlapping aspects of selection in modern populations. Number of children is a proxy for realized reproductive success, but in contemporary cohorts it is strongly influenced by cultural, socioeconomic, and behavioral factors (e.g., contraception)^78^ that may rapidly change over generations and hence reflect only recent and short-term evolutionary pressures. In contrast, parental longevity provides an indirect measure of survival-related fitness across generations, whereas the composite health score aggregates deviations across multiple physiological systems within the current generation and may therefore represent a broader measure of somatic maintenance. An important difference emerged between approaches leveraging rare *vs* common variants. Rare-variant burden estimates exhibited several consistent and biologically coherent relationships with multiple independent evolutionary constraint metrics, including SNP heritability, pLI, transcript evolutionary rate, and Moran’s I, although some associations varied across filtering thresholds. In contrast, common-variant-based selection estimates showed weaker and less consistent associations with orthogonal constraint measures. This difference may reflect the distinct genetic architectures and selective pressures captured by the two approaches. Rare loss-of-function variants are typically subject to stronger purifying selection and are disproportionately depleted within dosage-sensitive and functionally essential genes. Consequently, burden-based estimates may more directly reflect long-term gene-level functional constraint. By contrast, common-variant architectures inferred through LDpred2 may partly capture indirect, polygenic effects. Despite these differences, concordance analyses revealed some meaningful overlap across methods. Observational quadratic coefficients derived from the three fitness proxies showed modest positive correlations with one another, indicating that proteins influencing one component of fitness often also influence others, despite the fact that the three fitness proxies are not correlated with each other phenotypically (Supplementary Figure S4). Burden-based estimates showed strong agreement with selection signals derived from the health score. Rare loss-of-function variants disproportionately affect essential and dosage-sensitive genes, making it plausible that proteins under strong rare-variant constraint would also show measurable consequences for overall health. At the same time, the limited overlap observed across methods suggests that each framework captures partly distinct dimensions of selective constraint operating over different timescales. Our findings extend previous investigations of stabilizing selection on complex traits to circulating protein abundance, highlighting the potential relevance of selective constraint at the molecular level. Several limitations should nevertheless be considered. First, although we integrated multiple complementary approaches, all analyses remain fundamentally indirect and correlational. Second, our analyses primarily capture short-term microevolutionary signatures within contemporary human populations. Robust macroevolutionary analyses of protein abundance remain difficult because comparable cross-species proteomic datasets are not yet available at sufficient scale and depth. Third, the plasma abundance of 2’923 proteins represents only a fraction of the human proteome. Protein abundance is highly context-dependent and often varies substantially across tissues, cell types, and physiological states^14,79^. Our analyses were therefore limited to circulating proteins, which capture systemic molecular variation in blood but may not reflect tissue- or cell-type-specific regulation. Nevertheless, circulating proteins represent clinically important molecular phenotypes, and many exhibit substantial genetic regulation. Fourth, all analyses were restricted to individuals of European ancestry, potentially limiting generalizability to other populations with distinct demographic histories, allele-frequency spectra, linkage disequilibrium patterns, and environmental exposures. A fifth limitation arises from a fundamental challenge of inferring selection from genomic variation: the strongest stabilizing selection can erase the very genetic variation needed to detect it. Large-effect variants are rapidly driven to extremely low frequencies or removed entirely from the population, thereby reducing power to characterize frequency-dependent genetic architectures^39,2^. Future studies leveraging larger multi-ancestry proteogenomic datasets, longitudinal cohorts, and cross-species comparative proteomic resources will be essential to improve both statistical resolution and causal interpretability. The rapidly expanding availability of large-scale proteomic resources may ultimately enable a more comprehensive characterization of evolutionary constraints across tissues, populations, timescales, and multiple layers of molecular regulation.

## 5 Data and code availability

This study used data from the UK Biobank under application number 16389. Individual-level genetic, phenotypic, and proteomic data are available to approved researchers through the UK Biobank, subject to its access procedures and restrictions. UKBB data analyses were carried out in part in the High-Performance Computing Center of University of Lausanne, Switzerland. Code is available at https://github.com/dianazanoaga/ stabilizing_selection_human_proteins. Analyses were performed in R (v4.5.3). Polygenic modelling used bigsnpr (LDpred2-auto) together with Matrix and arrow; LASSO regressions used glmnet; dN/dS was estimated with orthologr, biomartr, Biostrings, metablastr, rdiamond and ape; transcript evolutionary rates were estimated with geiger and Moran’s I with spdep. Data handling used data.table, dplyr, tidyr, tibble, purrr, stringr, magrittr, readr, readxl and writexl, and figures were produced with ggplot2, patchwork, gridExtra, grid, ggrepel, ggpubr, scales and UpSetR.

## 6 Acknowledgments

The work was funded by the Swiss National Science Foundation (315230_2_19587).

## 8 Declaration of generative AI and AI-assisted technologies in the manuscript preparation process

During the preparation of this work, the author(s) used OpenAI and Claude to assist with language editing, improving the clarity and readability of the text, and developing and refining code for data analysis and visualization. The author(s) reviewed and edited the output as needed and take full responsibility for the content of the published article.

## 9 Supplementary

**Supplementary Figure S1:**
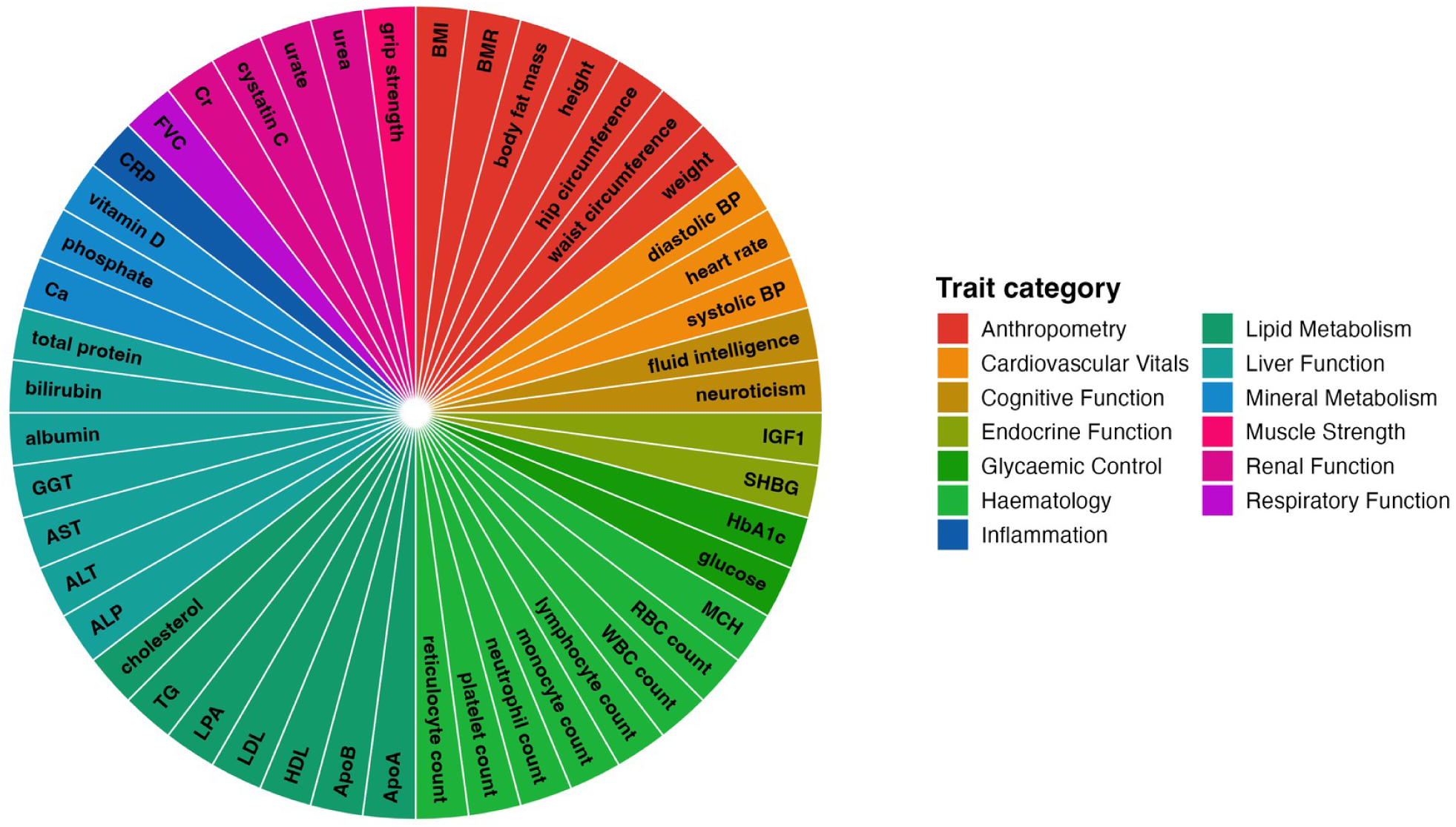
Overview of the 48 complex traits used for building the health score fitness proxy. Traits are grouped into biological categories (legend) and represent anthropometric, cardiovascular, metabolic, endocrine, cognitive, immune, liver, renal, pulmonary, and hematological phenotypes available in UK Biobank. These complex traits were used as potential mediators of protein–fitness selection signals.

**Supplementary Figure S2:**
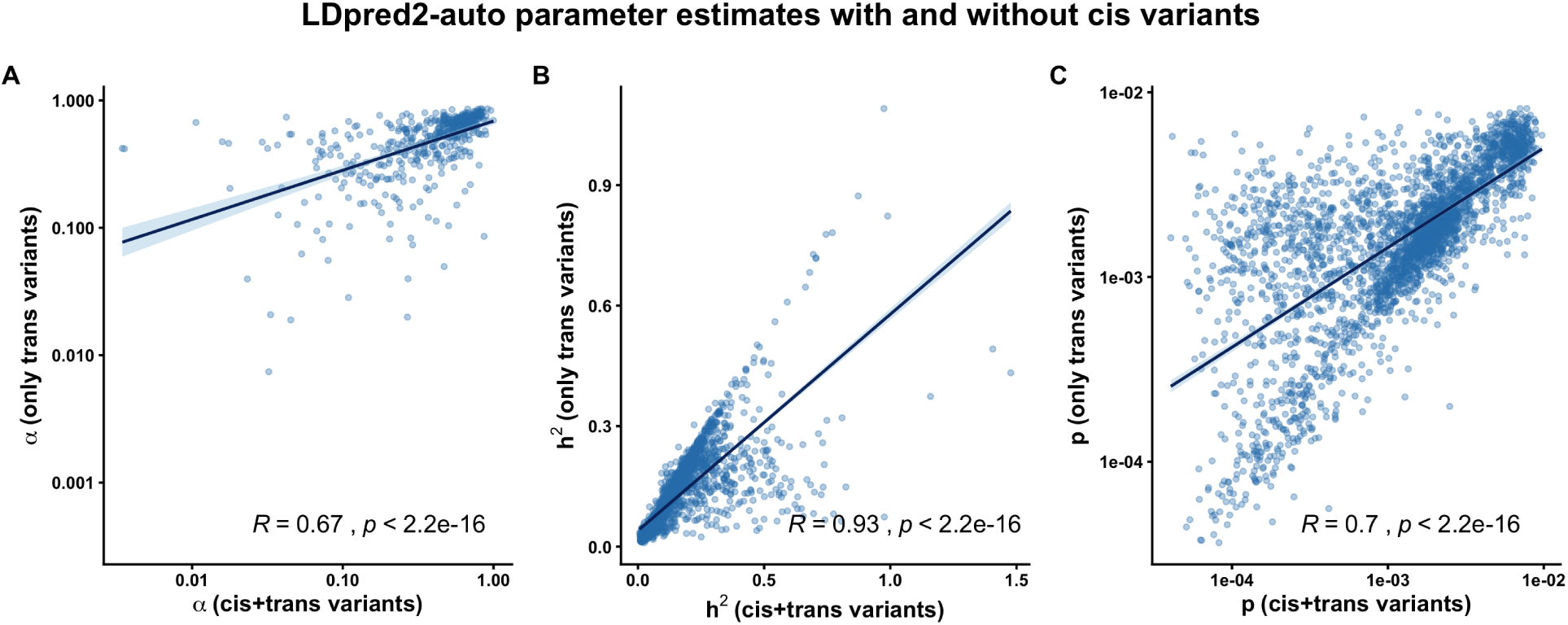
Concordance of LDpred2-auto parameter estimates using all variants (cis+trans) versus trans-only variants. Comparison of LDpred2-auto estimates for (A) the selection parameter *α*, (B) SNP-heritability (*h*^2^), and (C) the fraction of causal variants (*p*) obtained when fitting the model using all genome-wide variants (cis+trans) versus restricting to trans-acting variants only after excluding cis-pQTLs. Spearman correlations across analyses were *ρ* = 0.93 (*p <* 2.2 *×* 10*^−^*^16^) for SNP-heritability, *ρ* = 0.67 (*p <* 2.2 *×* 10*^−^*^16^) for *α*, and *ρ* = 0.71 (*p <* 2.2 *×* 10*^−^*^16^) for polygenicity (*p*). These results indicate that excluding cis variants does not alter the inferred genetic architecture or selection parameter estimates.

**Supplementary Figure S3:**
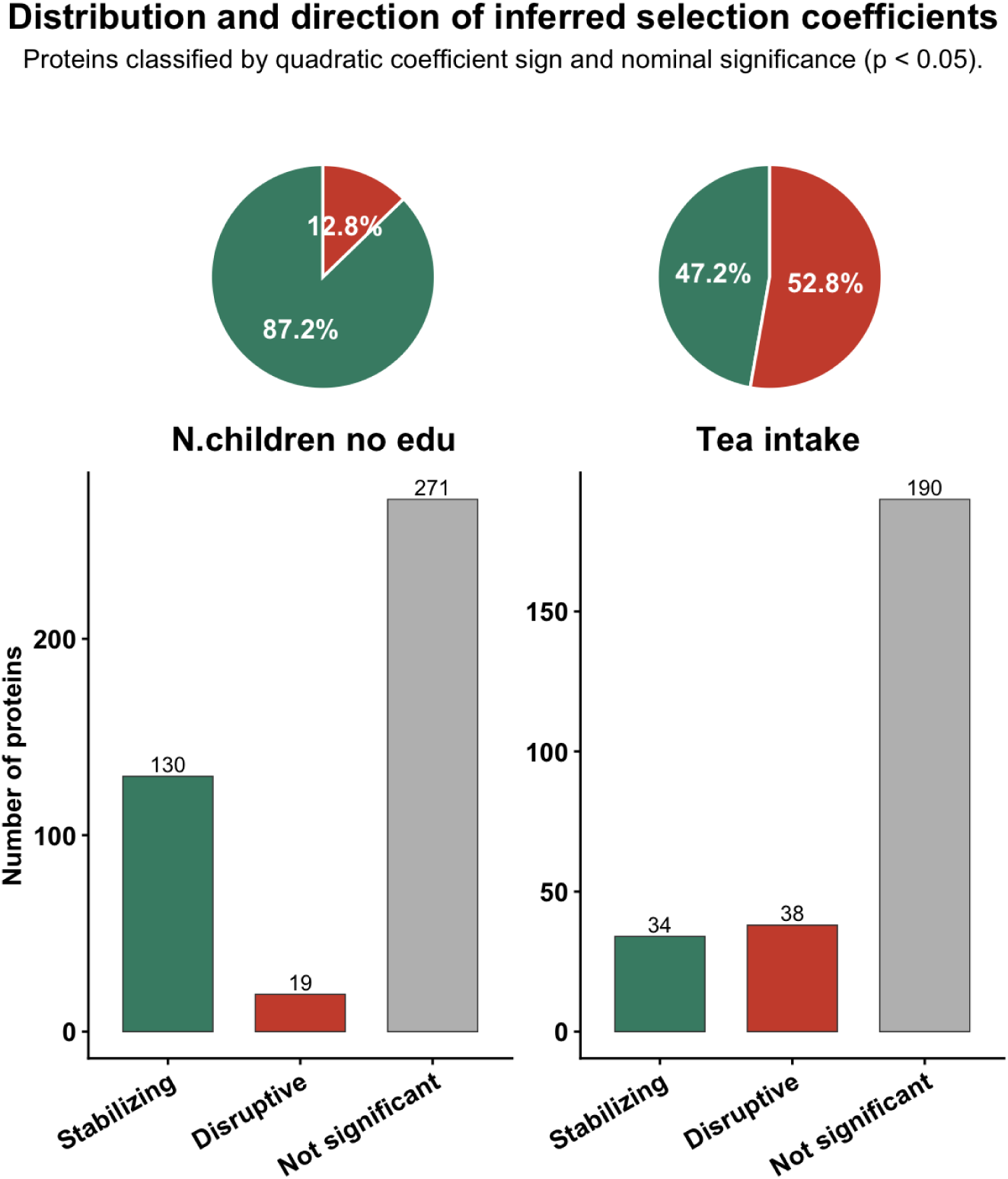
Distribution and direction of inferred selection coefficients for protein abundance using number of children (without education adjustment) and tea intake as a negative control. Proteins were classified according to the sign and significance of inferred quadratic selection coefficients into three categories: significant negative coefficients (*α <* 0, nominal *p <* 0.05), significant positive coefficients (*α >* 0, nominal *p <* 0.05), and non-significant proteins. Counts are shown for regression models relating observed protein abundance to two phenotypes: number of children estimated without adjustment for educational attainment, and tea intake included as a negative control phenotype. Pie charts above each panel show the relative proportion of significant negative versus significant positive coefficients among proteins reaching nominal significance. Only proteins with inferred optimum 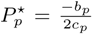 within *±*1 standard deviation of the observed protein distribution were included. The number of children analysis showed a predominance of significant negative coefficients, consistent with stabilizing selection signatures, whereas the tea intake analysis served as a negative control to evaluate whether similar patterns arise for a phenotype not expected to be directly related to fitness.

**Supplementary Figure S4:**
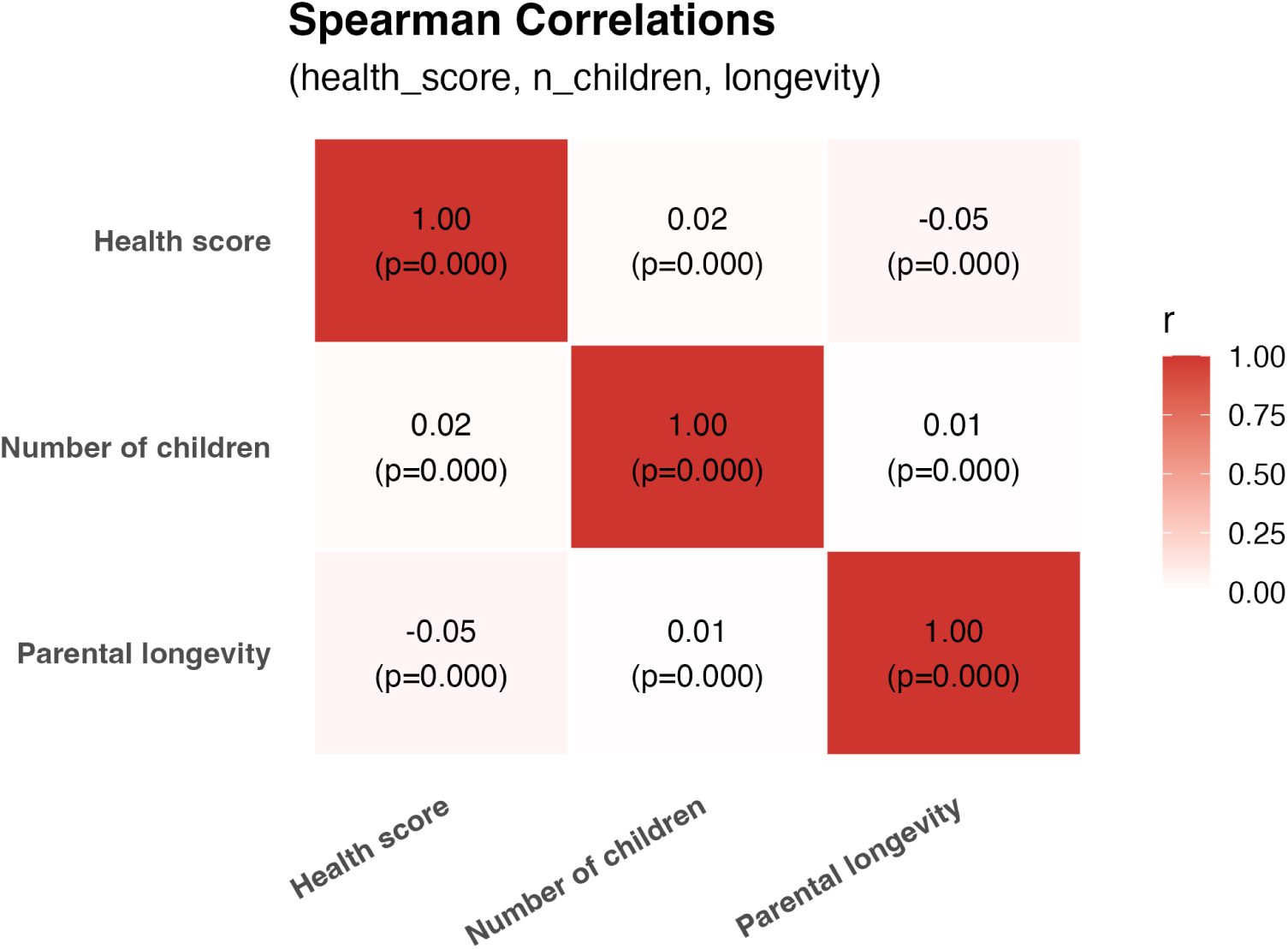
Fitness proxies are essentially uncorrelated. Spearman correlations among the three proxies (number of children, composite health score, and parental longevity) are near zero (all *ρ ≈* 0), indicating that these measures capture largely orthogonal axes of variation. Cells display *ρ* (top) and the corresponding *p*-value (bottom); color encodes *|ρ|*.

**Supplementary Figure S5:**
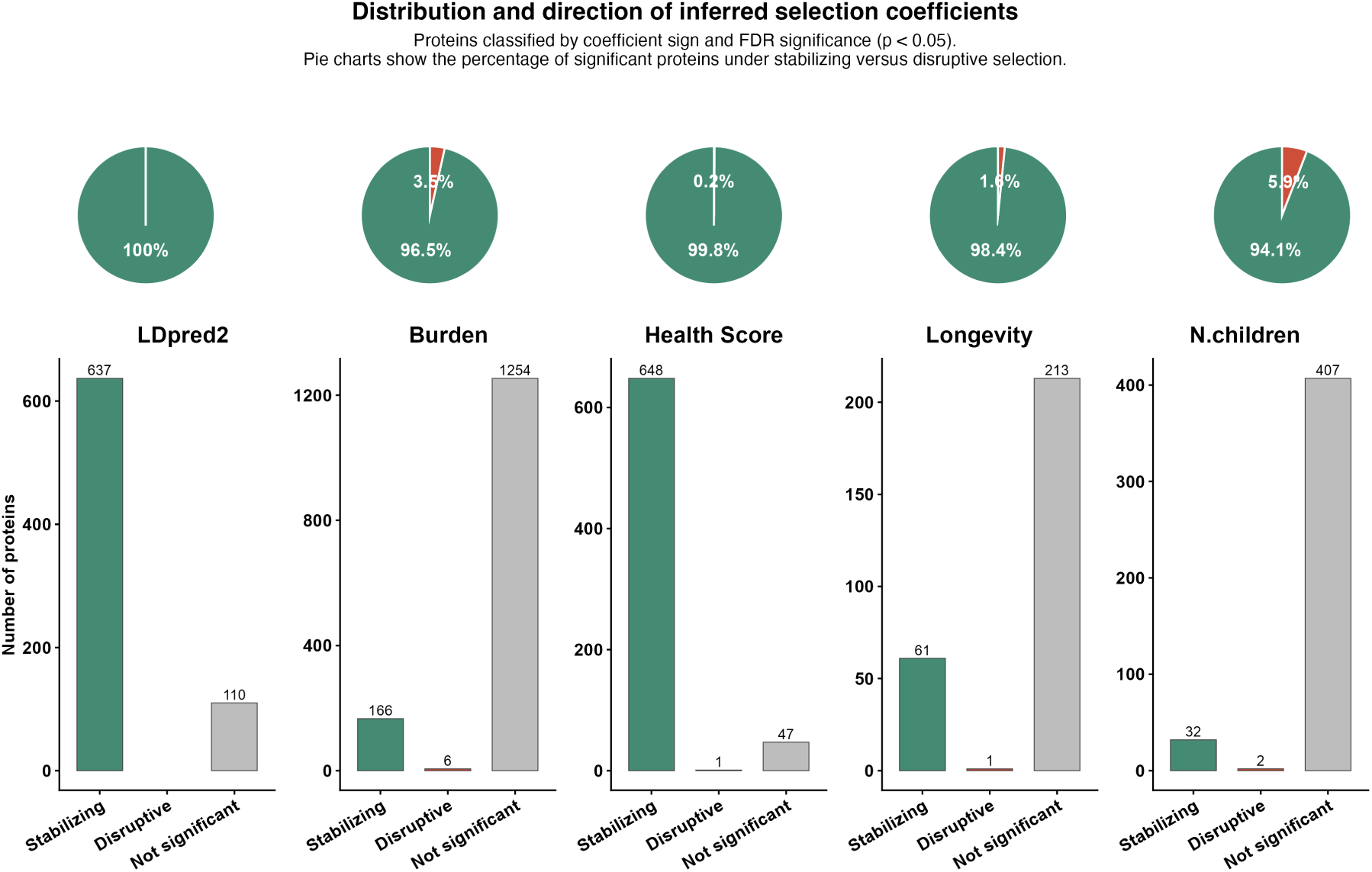
Distribution and direction of inferred selection coefficients across methods using FDR-adjusted significance thresholds. Proteins were classified according to the sign and significance of inferred selection coefficients into three categories: significant negative coefficients (*α <* 0, FDR-adjusted *p <* 0.05), significant positive coefficients (*α >* 0, FDR-adjusted *p <* 0.05), and non-significant proteins. Counts are shown for LDpred2-based estimates, rare variant burden analyses, and regression models relating observed protein abundance to three fitness proxies (health score, parental longevity, and number of children). Pie charts above each panel show the relative proportion of significant negative versus significant positive coefficients among proteins reaching FDR significance. For the regression analyses, only proteins with inferred optimum 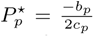 within *±*1 standard deviation of the observed protein distribution were included. Across all analyses, significant negative coefficients substantially outnumbered significant positive coefficients. Negative coefficients accounted for 94.1% to 100% of significant associations, indicating a strong predominance of stabilizing selection signatures across methods.

**Supplementary Figure S6:**
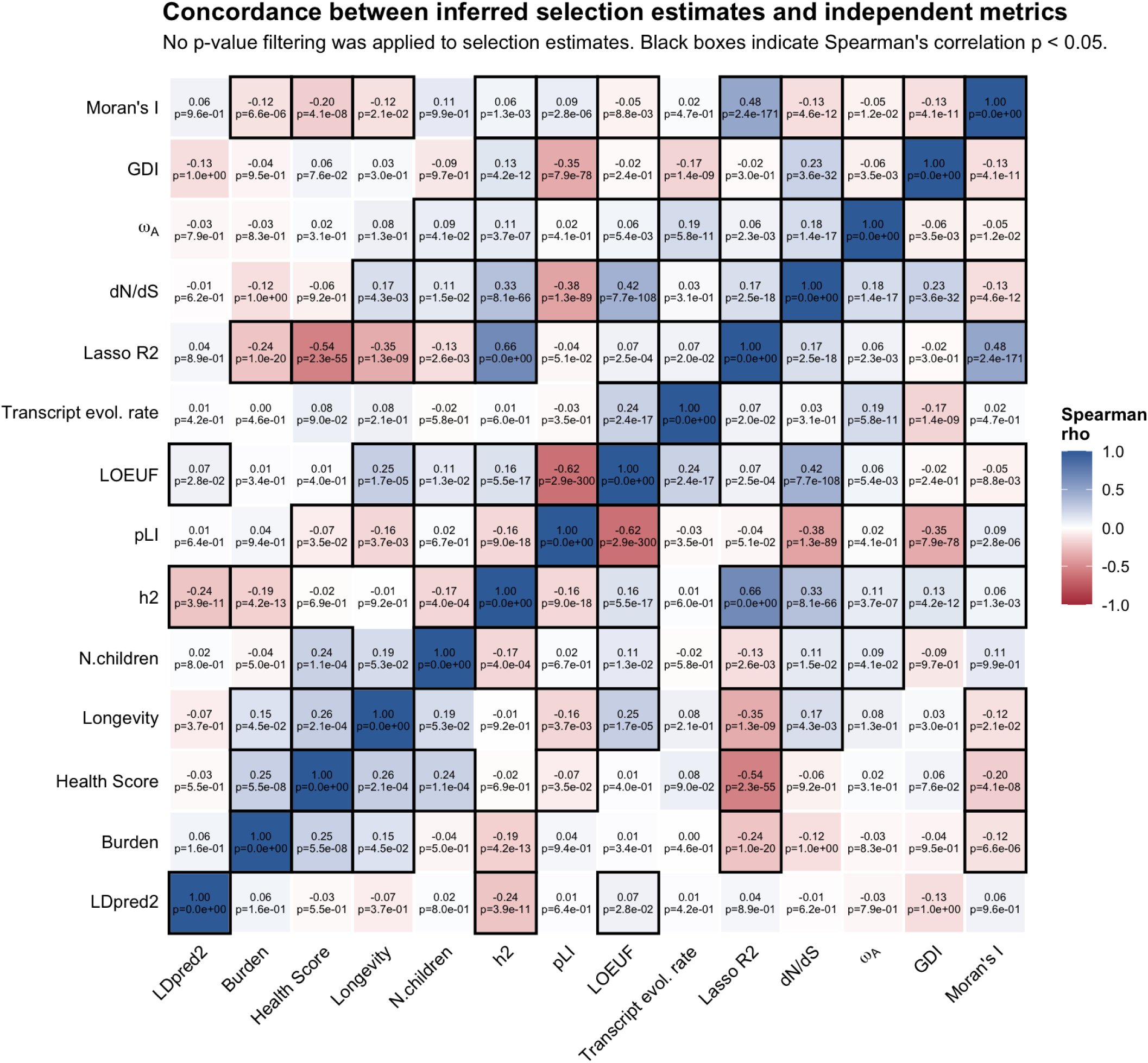
Concordance between inferred protein-level selection coefficients and independent metrics. Heatmap showing Spearman correlations between inferred selection coefficients and orthogonal measures of evolutionary and functional constraint, including protein heritability (*h*^2^), probability of loss-of-function intolerance (pLI), LOEUF, meta-analysed transcript evolutionary rate, LASSO *R*^2^, dN/dS, *ωA*, Gene Damage Index (GDI), and Moran’s I. Prior to correlation analyses, method-specific filtering criteria were applied to retain reliable estimates only. The heatmap reports correlations computed using inferred selection coefficients without applying any p-value filtering. Cell colors indicate Spearman correlation coefficients (*ρ*), and black borders denote statistically significant correlations (*p <* 0.05). Supplementary Figure S8 reports the number of overlapping proteins used for each pairwise comparison after applying all filtering criteria and significance thresholds.

**Supplementary Figure S7:**
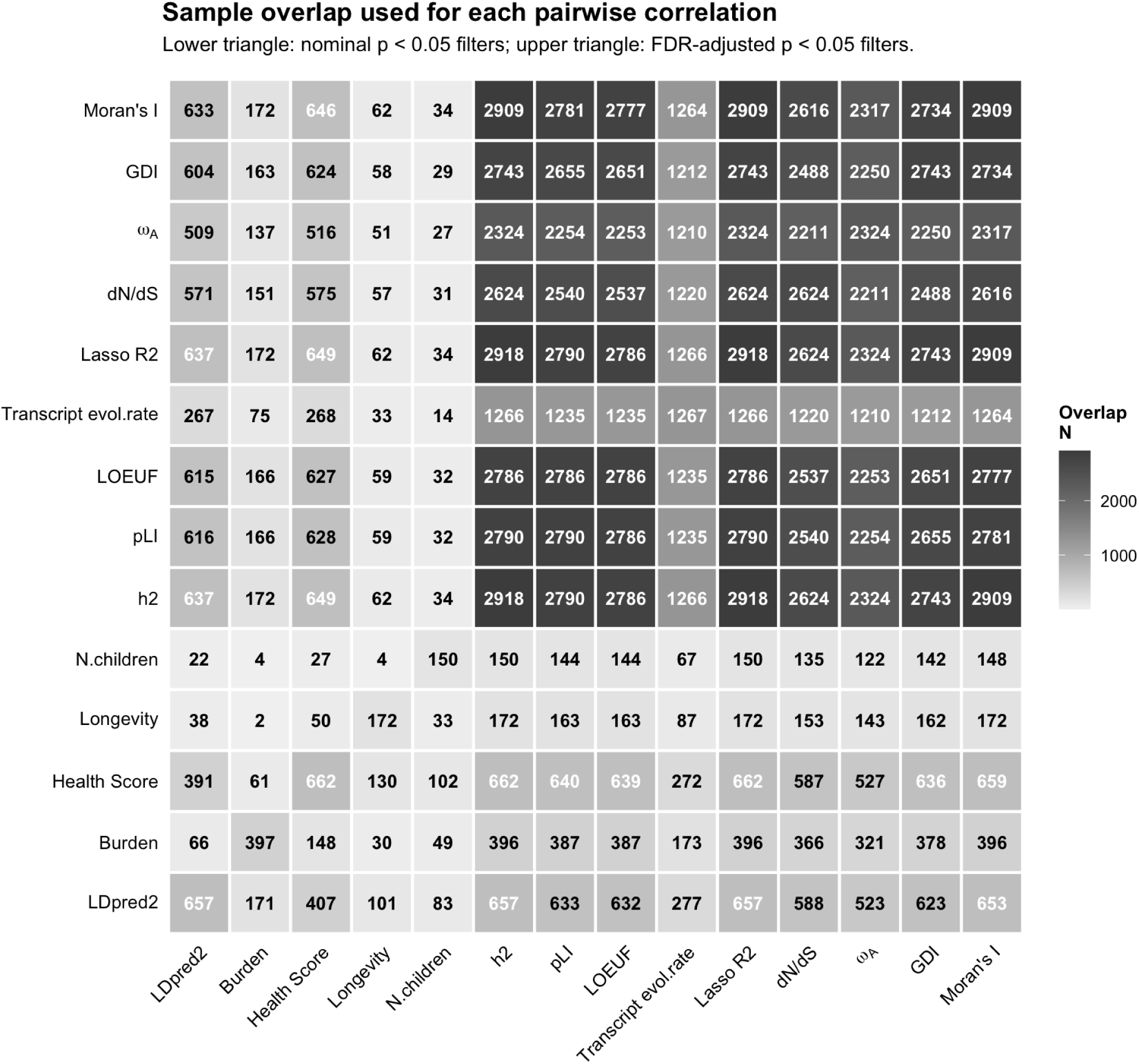
Number of overlapping proteins used for pairwise correlation analyses across inferred selection coefficients and independent metrics under nominal and FDR significance thresholds. Heatmap reporting the number of overlapping proteins used to compute each pairwise Spearman correlation shown in Figure 4 after applying all method-specific filtering criteria. The lower triangle corresponds to analyses restricted using nominal significance thresholds applied to inferred selection coefficients (*p <* 0.05), whereas the upper triangle corresponds to analyses restricted using false discovery rate corrected thresholds (*p <* 0.05). Values therefore represent the effective number of proteins jointly retained for each pairwise comparison after harmonization across datasets and implementation of all quality-control and significance-filtering procedures.

**Supplementary Figure S8:**
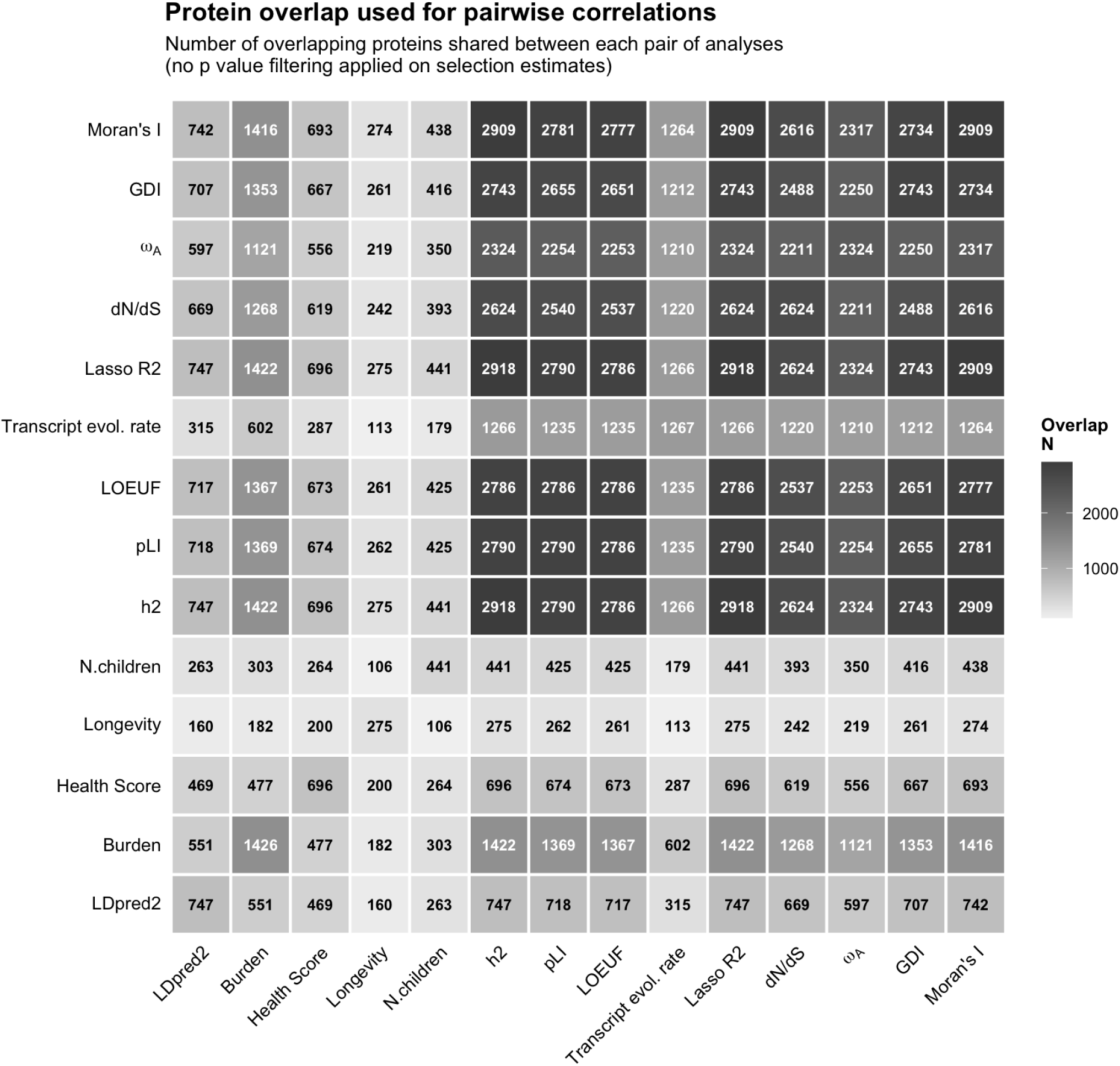
Number of overlapping proteins used for pairwise correlation analyses across inferred selection coefficients and independent metrics without significance filtering on selection estimates. Heatmap reporting the number of overlapping proteins used to compute each pairwise Spearman correlation shown in Figure S6 after applying all method-specific filtering criteria, but without imposing significance thresholds on inferred selection coefficients (*p <* 1). Values therefore reflect the effective number of proteins jointly available for each pairwise comparison following harmonization across datasets and implementation of all quality-control procedures independently of statistical significance filtering.

**Supplementary Figure S9:**
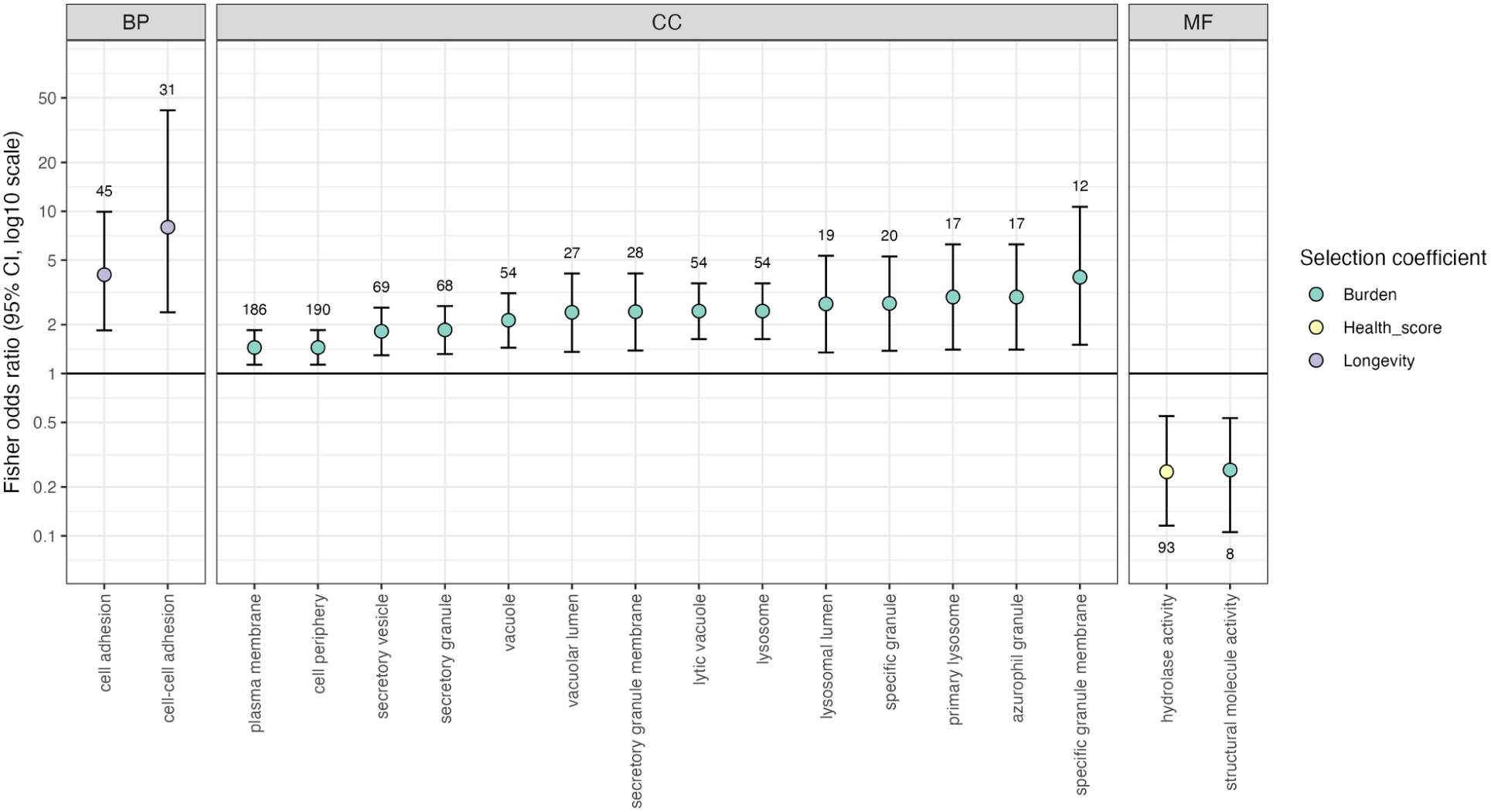
Gene Ontology enrichment analysis of proteins exhibiting stabilizing selection signatures. The figure shows the log10 odds ratios derived from Fisher’s exact tests. Enrichment tests were calculated separately for the three GO aspects (BP – biological process, CC – cellular component, MF – molecular function) and are shown in separate panels. The colors of the dots indicate based on which selection coefficient the given protein group was estimated to be under stabilizing selection, with the numbers next to each dot showing the number of proteins that are both associated with the given GO term and are under significant stabilizing selection. Only significantly enriched/depleted GO terms are shown, with a significance cutoff of FDR*<* 0.05.

